# Estimating residence times of lymphocytes in ovine lymph nodes

**DOI:** 10.1101/513176

**Authors:** Margaret M. McDaniel, Vitaly V. Ganusov

## Abstract

The ability of lymphocytes to recirculate between blood and secondary lymphoid tissues such as lymph nodes (LNs) and spleen is well established. Sheep have been used as an experimental system to study lymphocyte recirculation for decades and multiple studies exist documenting accumulation and loss of intravenously (i.v.) transferred lymphocytes in efferent lymph of various ovine LNs. Yet, surprisingly little work has been done to accurately quantify the dynamics of lymphocyte exit from the LNs and to estimate the average residence times of lymphocytes in ovine LNs. In this work we developed a series of mathematical models based on fundamental principles of lymphocyte recirculation in the body and specifically, on how lymphocytes enter and exit lymph nodes in non-inflammatory (resting) conditions. We fitted these models to data from several independent experiments. Our analysis suggested that in sheep, recirculating lymphocytes spend on average 3 hours in the spleen and 20 hours in skin or gut-draining LNs with a distribution of residence times in LNs following a skewed gamma (lognormal-like) distribution. The latter result is in contrast which recent suggestions that the distribution of residence times of naive T cells in murine LNs is exponential and that lymphocyte residence times depend on the LN type (e.g., gut- vs. skin-draining). Our mathematical models also suggested an explanation for a puzzling observation of the long-term persistence of i.v. transferred lymphocytes in the efferent lymph of the prescapular LN (pLN); the model predicted that this is a natural consequence of long-term persistence of the transferred lymphocytes in circulation. We also found that lymphocytes isolated from the skin-draining pLN have a two-fold increased entry rate into the pLN as opposed to the mesenteric (gut-draining) LN (mLN). Likewise, lymphocytes from mLN had a three-fold increased entry rate into the mLN as opposed to entry rate into pLN. Importantly, these cannulation data could not be explained by preferential retention of cells in LNs of origin. Taken together, our work illustrates the power of mathematical modeling in describing the kinetics of lymphocyte migration in sheep and provides quantitative estimates of lymphocyte residence times in ovine LNs.

## 1 Introduction

One of the peculiar properties of the adaptive immune system of mammals is the ability of its cells, lymphocytes, to recirculate between multiple tissues in the body; that is lymphocytes in the blood are able to enter the tissues and after some residence times in the tissues, they return to circulation [1]. The pattern of lymphocyte recirculation in general depends on the lymphocyte type (e.g., B or T cell), status of the lymphocyte (resting vs. activated), and perhaps tissues via which lymphocytes are migrating. Naive, antigen-unexperienced lymphocytes, primarily recirculate between secondary lymphoid tissues such as lymph nodes, spleen, and Peyer’s patches [2–4]. Following activation after exposure to an antigen, naive lymphocytes become activated and differentiate into effector lymphocytes which are able to access non-lymphoid tissues such as the skin and gut epithelium [2, 3, 5–7].

Why lymphocytes recirculate is not entirely clear [8, 9]. Because the frequency of lymphocytes specific to any given antigen is in general low (e.g., [10]) and the place of entry of any pathogen is unknown by the naive host, recirculation of lymphocytes may increase the chance to encounter the infection by rare pathogen-specific lymphocytes [11]. However, evidence that impairing lymphocyte recirculation influences ability of the host to respond to infections is very limited. For example, the use of the drug FTY720 (fingolimod) in humans has been associated with a higher incidence of severe infections [12, 13]. FTY720 is believed to reduce the ability of lymphocytes to exit lymph nodes, thus, reducing their ability to recirculate [14–16]. However, whether the side-effects of fingolimod is exclusively due to its impact on recirculation ability of lymphocytes in humans is unknown.

The ability of lymphocytes to recalculate between blood and lymph have been nicely demonstrated in now classical experiments by Gowans in rats and later by Hall and Morris in sheep [17, 18]. Interestingly, subpopulations of lymphocytes can migrate preferentially to different regions of the body, based on their origin as well as their type [19–24]. Molecular interactions between receptors and associated ligands corresponding to the selective entry of lymphocytes to both lymphoid and non-lymphoid tissue have been relatively well-characterized [25–28]. However, the actual kinetics of lymphocyte recirculation have been characterized mostly qualitatively, and we still do not fully understand how long lymphocytes reside in the spleen, LNs, and Peyer’s patches, and how such residence times depend on lymphocyte type and animal species, especially for humans.

Understanding lymphocyte migration via lymph nodes (LNs) may be of particular importance for larger animals, including humans, where LNs constitute the majority of the secondary lymphoid tissues [29, 30]. Lymphocytes may enter the LNs via two routes: from the blood via high endothelial venules (HEVs) or from the afferent lymph draining interstitial fluids from surrounding tissues [1]. Experimental measurements suggest that under resting, noninflammatory conditions most lymphocytes (about 80 −90%) enter the LNs via HEVs [31]. Lymphocytes in LNs exit the nodes with efferent lymphatics which either passes to the next LN in the chain of LNs or via right or left lymphatic ducts return to circulation [1, 9, 32]. In contrast, cells can only enter the spleen from the blood and cells exiting the spleen directly return to circulation.

In the past, to study lymphocyte recirculation via individual LNs sheep or cattle have been used [18, 33– 38]. In such experiments, lymphocytes are collected from specific tissue of the animal, e.g., blood, a removed LN, or efferent lymph of a LN, labeled with a radioactive or fluorescent label, and then re-infused back into the same animal (e.g., intravenously, i.v.). The dynamics of the labeled cells could then be monitored in the blood, or more commonly, in the efferent lymph of different LNs over time [35, e.g., see Figure 1A]. This is done by inserting a cannula and collecting lymph and cells exiting the LN over time. The dynamics of the labeled cells in the efferent lymph of various LNs follow a nearly universal pattern — the number of labeled cells increases initially, reaches a peak and then slowly declines over time [20, 31, 39, 40, and see Figure 2C&D]. Given that in many such experiments, the peak of labeled cells in the efferent lymph is reached in 24 hours and declines slowly, from the asymmetry of this time course (e.g., Figure 2C) it can be interpreted that the average residence time of lymphocytes in the ovine lymph nodes is about 48 hours. To our knowledge, the actual residence time in ovine LNs has not been regularly reported. Interestingly, with the use of mathematical modeling an accurate quantification of how long lymphocytes spend in LNs in sheep has been recently performed [41].

**Figure 1:**
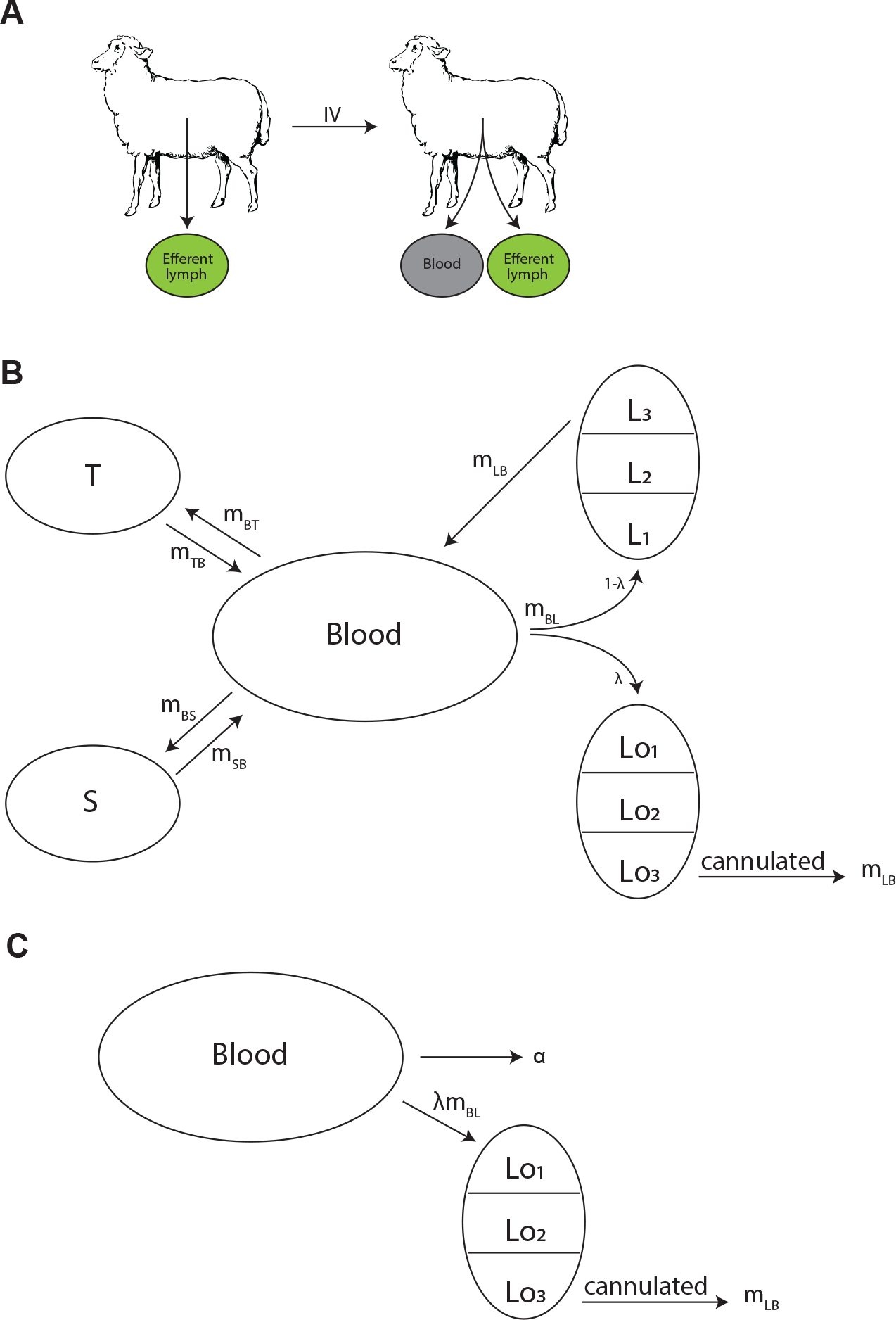
Experimental design and schematics of mathematical models on lymphocyte migration in sheep. In experiments, recirculating lymphocytes were obtained from efferent lymph of various LNs, labeled in vitro, and then re-injected i.v. into the same animal. The concentration of injected lymphocytes was then followed in sheep in the blood and the efferent lymph of a given LN (panel A). To describe migration kinetics of recirculating lymphocytes via LNs we adopt a basic mathematical model from our previous study [43]. In the “recirculation” model (panel B) lymphocytes in the blood *B* may migrate to three (*n* = 3) major tissue compartments: spleen *S* (at a rate *m*_*BS*_), lymph nodes *L* (at a rate *m*_*BL*_), or other peripheral tissues *T* (at a rate *m*_*BT*_). A fraction *λ* of cells migrating to lymph nodes migrate to the cannulated LN (*Lo*) and thus can be measured, and remaining cells migrate to other lymph nodes (*L*). Exit of lymphocytes from the spleen or peripheral tissues follows first order kinetics at rates *m_SB_* and *m_TB_*, respectively (and thus residency times in these tissues are exponentially distributed). In contrast, following our previous work [43] residency times in LNs are gamma distributed modeled by assuming *k* sub-compartments (in the figure *k* = 3); exit from each sub-compartment is given by the rate *m*_*LB*_. In the alternative, ‘blood-LN dynamics’ model (panel C) cells in the blood (*B*) enter the cannulated LN at a rate *λm*_*BL*_ and exit the LN by passing via *k* sub-compartments at a rate *m*_*LB*_. Cells in the blood also leave the blood at a rate *α* (in case of a single exponential decline).

**Figure 2:**
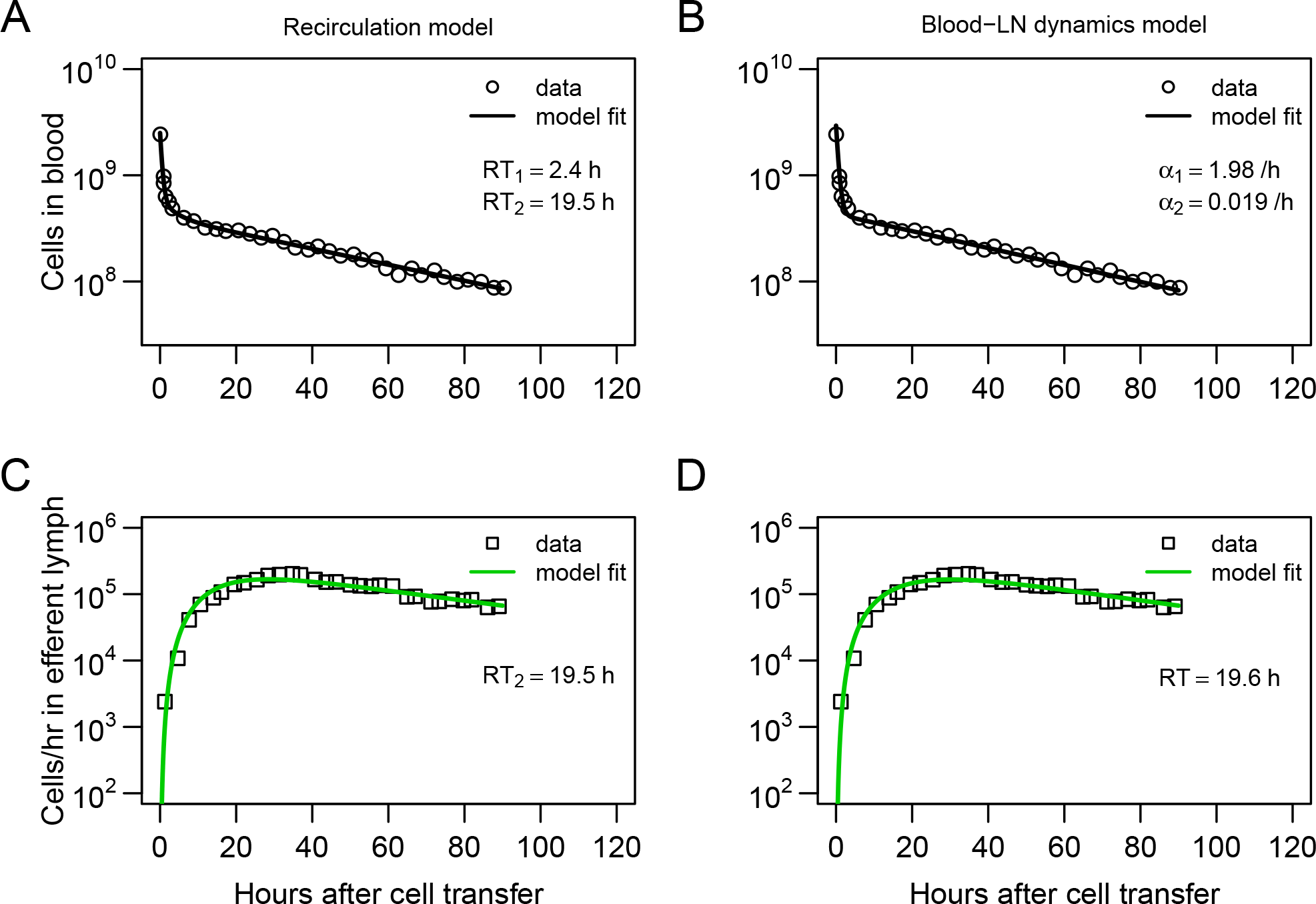
Dynamics of recirculating lymphocytes (RLs) in the blood naturally explains kinetics of accumulation and loss of RLs in prescapular LNs (pLN) of sheep. RLs were collected by cannulating pLN in sheep and then reinfused into the animal (see Frost *et al.* [39], Materials and Methods and Figure 1 for more detail). Dynamics of RLs in the blood (panels A&B) and in the efferent lymph of the pLN (panels C&D) for the first 90 hours after cell transfer (short-term migration) are shown by markers. We fitted two alternative models to these data. The first model assumed that the lymphocytes in the blood are able to recirculate between several tissue compartments such as spleen, LNs, and other tissues (see Figure 1B and eqns. (3)–(9)); the model with *n* = 3 tissue compartments (spleen, LNs, and other non-lymphoid tissues) and *k* = 3 sub-compartments in the LNs was able to accurately explain the data (panels A&C). The average residence time estimated for the two first compartments (*RT*_1_ and *RT*_2_) are indicated in the panels. The second, blood-LN dynamics model considers that the loss of RLs from the blood is described by phenomenologically by *j* = 2 exponential functions (with rates *α*_1_ and *α*_2_) and that LNs have *k* = 3 sub-compartments (see Figure 1C and eqns. (10)–(12)). The estimated average residence time of lymphocytes in the pLN is indicated in panel D as *RT*. Fits of models that assume different numbers of tissue compartments, different number of sub-compartments in LNs, or different numbers of exponential functions are shown in Tables 1 and S1. Parameters for the best fits of these models are given in Table 2.

In their pioneering study Thomas *et al.* [41] analyzed data on migration of lymphocytes via individual ovine LNs. To quantify these dynamics, the authors developed a mathematical model which considers cell migration via a LN as a random walk between multiple sub-compartments in the LN. In the model cells entering the LN start in the first sub-compartment, progress via the series of subcompartments by a random walk and eventually exit the node by leaving the last subcompartment [41]. As their model allows for the possibility for cells to spend variable lengths of time in the LN, it can naturally explain the long duration of labeled lymphocytes exiting the LN. By fitting the model to several different sets of data, the authors concluded that the average residence time of lymphocytes in ovine LNs is about 31 hour [41].

In addition to the average residence time the distribution of residence times may be important. In particular, if lymphocytes that just entered the LN have the same chance of exiting it as lymphocytes that already spent some time in it (i.e., distribution of residence times is exponential), this could suggest that exit of lymphocytes from LN is a simple Poisson-like stochastic process. Indeed, one recent study suggested that residency time of naive CD4 and CD8 T cells in LNs of mice is exponentially distributed [42]. In contrast, if lymphocytes require some time to be spent in the LN, for example, to acquire the ability to exit the node, then the distribution of residence time cannot be exponential. Recent work from our group suggests that residence time of thoracic duct lymphocytes in LNs of rats cannot be exponential and is best described by a gamma distribution with the shape parameter *k* = 2 or 3 [43]. Our analysis of data on kinetics of lymphocyte exit from inguinal LNs of photoconvertable Kaede mice did not allow to firmly establish the shape of the residence time distribution [9]. Whether the distribution of residence time of lymphocytes in ovine LNs is exponential or more complex is unknown.

In this paper, we formulated a series of mathematical models aimed at describing the kinetics of recirculation of lymphocytes in sheep. All models take into account basic physiological constrains on lymphocyte migration, for example, that lymphocytes enter the LNs continuously from the blood and lymphocytes that exit LNs return back to circulation. The models were fitted to a series of experimental data from previously published studies on lymphocyte migration in sheep. Our results suggest that the distribution of residence times of lymphocytes in ovine LNs is best described by a non-exponential distribution with estimated average residence times being 12-22 hours which depends on the type of lymphocytes used in experiments. The long-term presence of labeled lymphocytes in efferent lymph of cannulated LNs was explained by a continuous entry of new cells from the blood to the LN, thus, simplifying a previous modeling result [41]. Overall, our analysis provides a quantitative framework to estimate kinetics of lymphocyte recirculation using measurements of lymphocyte numbers in the blood and efferent lymph of ovine LNs. Such a framework may be useful to understand the efficacy and potential limitations of immunotherapies involving adoptive transfer of T cells in humans [44–46].

## 2 Materials and methods

### 2.1 Experimental data

For our analyses we digitized the data from several publications [20, 39, 47]. We describe in short design of these experiments and how the data have been collected. For more detail the reader is referred to the original publications.

#### Lymphocyte dynamics in blood and efferent lymph (dataset #1)

Experiments of Frost *et al.* [39] have been performed with 4-24 month old Alpenschalf or Schwarzkopf sheep (Figure 1A). Lymphocytes were collected from the efferent lymph of different lymph nodes (e.g., prescapular) at room temperature in sterile bottles with heparin and penicillin-streptomycin then labeled with ^51^Cr for 30 minutes after being washed and resuspended in phosphate buffered saline. For i.v. infusions 131 and for taking blood samples, a surgical procedure to install an indwelling cannula into the jugular vein was performed. The efferent duct of specific lymph nodes was also cannulated to allow for collection of lymph that passed through the node. The number of labeled lymphocytes in efferent lymph was determined by washing cells in a phosphate buffered saline and counting with a gamma scintillation counter. Labeled lymphocytes in venous or peripheral blood were counted similarly by a gamma scintillation counter then compared to the activity of plasma from the same volume of blood.

^51^Cr labeled lymphocytes were injected i.v. and the efferent lymph of prescapular LN (pLN) was collected. Lymph was collected as described above at 20-minute intervals for 3 hours or daily for about two weeks. The number of labeled lymphocytes in the lymph node was expressed as cpm per 10^7^ cells collected. Peripheral blood samples were only collected for certain experiments and they were taken at 10 minutes, 30 minutes, 1 hour, and 3-hour intervals thereafter. The number of labeled lymphocytes in peripheral blood was expressed as cpm/ml of whole blood minus the activity in the plasma. Cells exiting the lymph node were measured for 120 hours, while cells in the blood were measured for 90 hours, then once more at the 120^*th*^ hour. For this reason, we discuss lymphocyte migration kinetics in terms of short- and long-term migration experiments. The short-term experiments include the dynamics of the labeled lymphocytes in both blood and efferent lymph for the first 90 hours, and the long-term dataset considers all available data (measurement up to 120 hours).

According to Blood Volume of Farm Animals, Hampshire sheep less than a year to 3 years old have on average 6.3-5.8 ml blood per 100 gram weight and an average weight of 92-156 lbs [48]. This results in about 2.63 L of blood for Hampshire sheep less than a year old and about 4.1 L for Hampshire sheep 2-3 years old. The breed of sheep used in this study weight from 60 kg (Alpenschaf) to 100 kg (Schwarzkopf) at maturity, so it is reasonable to assume that these total blood estimates are less than those calculated for Hampshire sheep. We assume the volume of blood in the sheep is an average of these given volumes, specifically *V* = 3.4 L. This estimate was used to convert the estimate of the number of labeled lymphocytes per ml of blood to the total number of lymphocytes in the whole blood.

In one set of experiments (Figure 1 in Frost *et al.* [39]) 2.5 × 10^9^ labeled cells (representing 12.6×10^6^ cpm) were injected i.v. into sheep corresponding on average *RL*_*o*_ = 200 cells/cpm. Because in the original data, the cells in the blood at time *t* (*RL*_*t*_) were measured in cpm/ml of blood, the total number of cells in the blood *B* was calculated using the formula

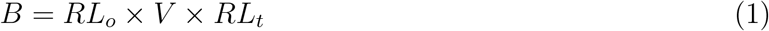

with *V* = 3.4 L and *RL*_*t*_ was digitized from Figure 1 of Frost *et al.* [39]. In the same experiments labeled cells exiting the pLN were measured as cpm/10^7^ cell with each data point summing 3 hours of cells collected every 20 minutes. Therefore, the amount of labeled cells exiting the pLN per hour (*C*_*t*_) is one third of this number. The total output of the pLN is given in Figure 3 in Frost *et al.* [39] and was estimated to be *f* = 10.85 × 10^7^ cell/hr. We calculate total cells exiting the pLN per hour by:

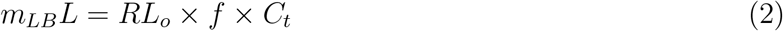

where *RL*_*o*_ = 200 cells/cpm. The final data (**`dataset1.csv`**, given as Supplement to the paper) includes changes in the total number of labeled lymphocytes in the peripheral blood and the number of labeled lymphocytes exiting pLN per hour over time.

**Figure 3:**
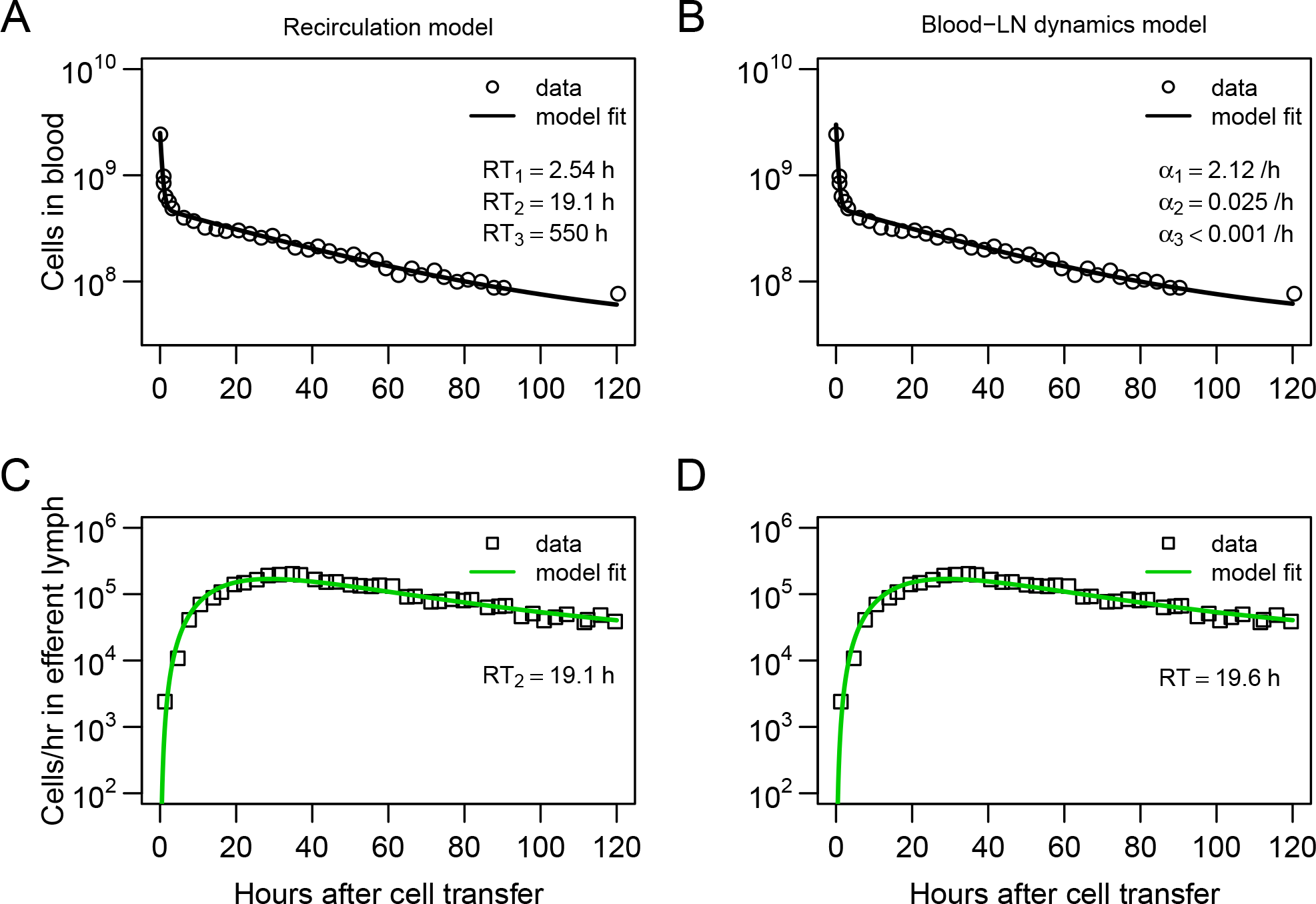
Explaining long-term recirculation kinetics of lymphocytes in sheep requires more complex mathematical models. We fitted either recirculation model (panels A&C, see eqns. (3)–(9)) or the blood-LN dynamics model (panels B&D, see eqns. (10)–(12)) to the data on the dynamics of labeled lymphocytes in the blood (panels A-B) and in efferent lymph of the pLN (panels C-D; see Materials and Methods for more detail) sampled for 5 days. In these experiments, there was only a single additional measurement of the labeled lymphocytes in the blood but continuous measurement of labeled lymphocytes in the efferent lymph (e.g., compare panel A to Figure 2A). The best fit of the recirculation model was found assuming lymphocyte recirculation via *n* = 3 different compartments (and *k* = 3 sub-compartments for the LNs) and average residency times in these compartments are indicated in panel A as *RT*_*i*_. The best fit of the blood-LN dynamics model was with *j* = 3 exponential decay functions and *k* = 3 sub-compartments for the LNs. The best fit models were determined via a series of trials (see Tables S1 and S2). Parameter estimates and 95% CIs are shown in Table 2.

#### Migration of lymphocytes from afferent to efferent lymph (dataset #2)

To further investigate how lymphocytes migrate via ovine LNs we digitized data shown in Figure 4 of Young *et al.* [47]. In these experiments different subsets of lymphocytes (CD4 T cells, CD8 T cells, *γδ* T cells, and B cells) were collected from efferent prescapular lymph and labeled with PKH-26 or CFSE to distinguish between different cell subsets. A maximum of 2×10^6^ cells of each type were infused into two popliteal afferent lymphatics over one hour. Cells were collected as they exited the cannulated efferent lymph of the popliteal LN (poLN), and phenotyped. The final data (**`dataset2.csv`**, given as Supplement to the paper) includes the percent of labeled cells found in the efferent lymph of the poLN over time.

**Figure 4:**
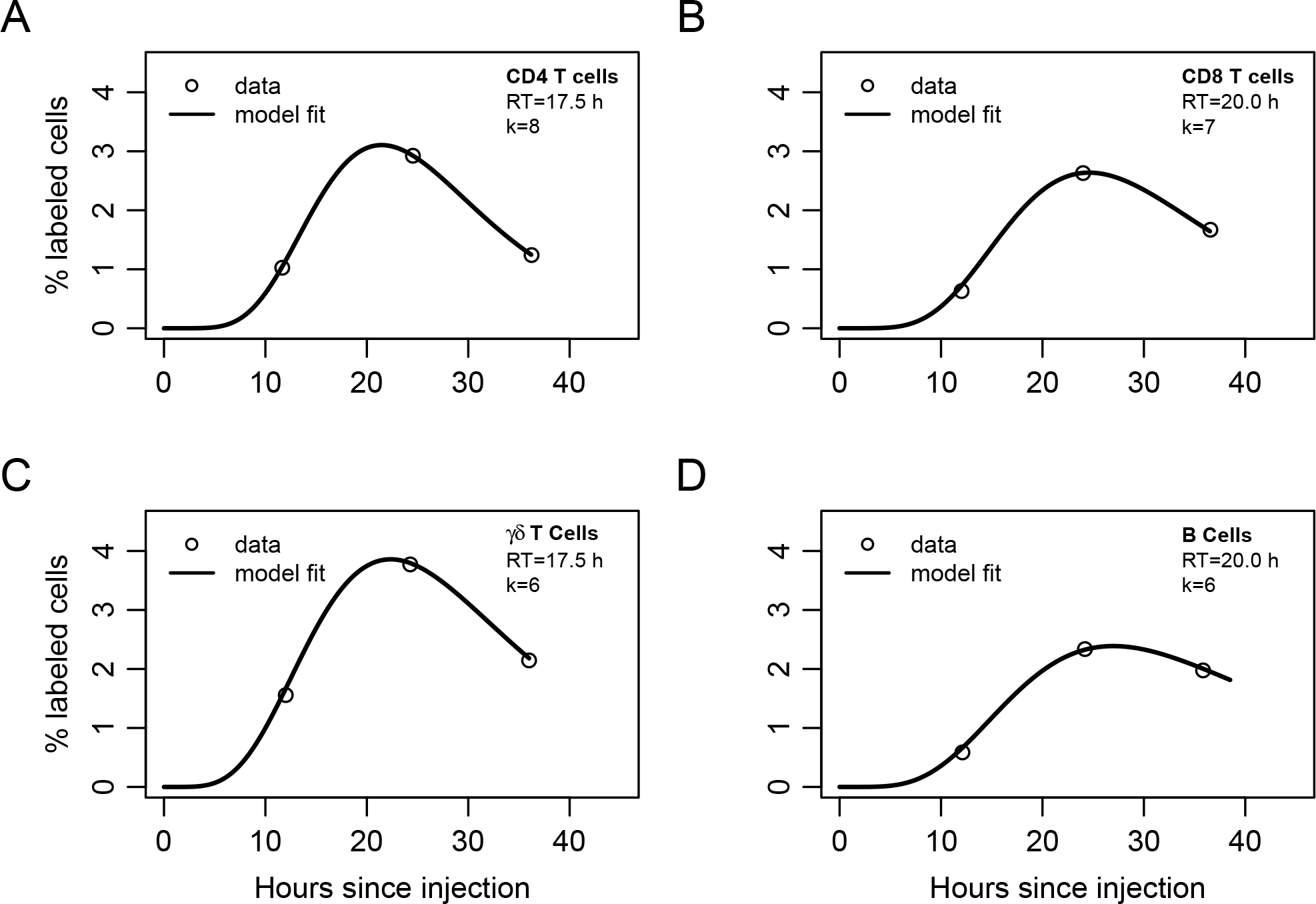
Mathematical modeling suggests non-exponentially distributed residency time of different subsets of lymphocytes in the LNs. CD4 T cells (panel A), CD8 T cells (panel B), *γδ* T cells (panel C) or B cells (panels D) were labeled and then injected into afferent lymph of popliteal LN (poLN) of sheep. The percent of labeled cells was followed in efferent lymph over time (see Materials and Methods for more detail and Young *et al.* [47]). We fitted a series of mathematical models assuming migration of injected lymphocytes into the LN with a variable number of sub-compartments *k* in the LNs (eqns. (13)–(15)). Due to scarcity of the data it was not possible to accurately estimate all parameters of the model (Table S4) and therefore the analysis was performed by assuming several fixed values for the residence time *RT* and by varying the number of sub-compartments in LNs (see Table S3). The best fits of the model leading to the lowest AIC values with the noted number of sub-compartments in the LN (*k*) are shown by lines and parameters of the models are given in Table S4.

#### Migration of T lymphocytes via skin-draining and gut-draining lymph nodes (dataset #3)

The final set of experimental data we used come from recirculation experiments of Reynolds *et al.* [20] with young sheep (Figure 6). Lymphocytes were isolated by cannulating the skin-draining pLN or the ileal end of the gut-draining mesenteric lymph node chain (mLN). Collected lymphocytes were passed through nylon wool columns to reduce the proportion of cells with surface immunoglobulin to less than 5% of the total, thus, creating T cell-enriched sample. Cells from intestinal or prescapular lymph were labeled with FITC or RITC, respectively. Labeled cells were re-infused into the same animal i.v. and cell density (given in labeled cells per 10^4^ cells) was reported for 240 hours (Figure 1 in Reynolds *et al.* [20]). Because the authors did not report the overall cell output in efferent lymph of pLN and mLN, we used the provided numbers of the frequency of labeled cells in in the lymph for fitting the models. Existing data suggest similar output of lymphocytes from pLN (1 − 5 × 10^8^ cells/h) or mLN (1 − 10 × 10^8^ cells/h, [35]) which in part justifies our approach. The final data (dataset3.csv, given as Supplement to the paper) includes the number of labeled cells from skin or intestinal lymph per 10^4^ cells found in efferent lymph of pLN or mLN over time during cannulation.

The raw data from the cited papers were extracted using digitizing software Engauge Digitizer (digitizer.sourceforge.net).

### 2.2 Mathematical Models

#### 2.2.1 Basic assumptions

In our models we assume that blood is the main supplier of lymphocytes to other tissues and that when exiting these tissues, lymphocytes will return to the blood [43, 49]. We also assume that all LNs behave similarly with respect to exit rates of lymphocytes from the node, and that cell infusion does not impact lymphocyte migration via individual LNs. The assumption that residency times of lymphocytes are similar in different LNs has been found in some but not all previous studies [42, 43].

#### 2.2.2 Models to predict lymphocyte dynamics in efferent lymph of LNs

##### Recirculation model

To predict the dynamics of i.v. transferred labeled lymphocytes in the blood and efferent lymph of pLN we extended a previously proposed compartmental model describing recirculation kinetics of lymphocytes in the whole body [43]. The model (Figure 1B) predicts the number of i.v. transferred lymphocytes in the blood (*B*), spleen (*S*), lymph nodes (*L*), and other non-lymphoid tissues (*T*). In the model we ignored migration of lymphocytes via vasculature of the lung and liver since previous work suggested that resting lymphocytes pass via these tissues, at least in rats, within a minute [43]. This is different from migration of activated lymphocytes via these tissues which could take hours [9]. In the model, cells in the blood can migrate to the spleen, lymph nodes, or other tissues at rates *m*_*Bi*_ and cells can return to circulation from these tissues at rates *m*_*iB*_ where *i* = *S, L, T*. When exiting spleen or non-lymphoid tissues lymphocyte follow the first order kinetics, so the decline of cells in the tissues in the absence of any input is given by an exponential function. This in part is based on our previous work suggesting of migration of thoracic duct lymphocytes via spleen can be described as first order kinetics [43]. In contrast, migration of lymphocytes via LNs may not follow the first order kinetics (e.g., [43]) and thus was modelled by assuming *k* sub-compartments in the nodes with equal transit rates *m*_*LB*_. Such subcompartments may represent different areas in the LNs, for example, paracortex and medulla. Mathematically, we used the sub-compartments to model non-exponential residency time of lymphocytes in the LNs.

To describe accumulation and loss of labeled lymphocytes in the cannulated lymph nodes we assume that a fraction of lymphocytes *λ* migrating to lymph nodes migrate to the cannulated node *Lo*_1_ (e.g., pLN, Figure 1B) while 1 − *λ* cells migrate to other LNs (*L*_1_). We assume that cells do not die but the process of migration to non-lymphoid tissues with no return back to circulation is equivalent to cell death. We did consider several alternative models in which death rate was added to the model (see Discussion). Taken together, with these assumptions the mathematical model for the kinetics of lymphocyte recirculation in ovine LNs is given by equations:

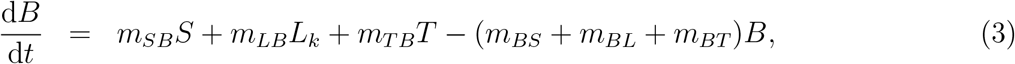

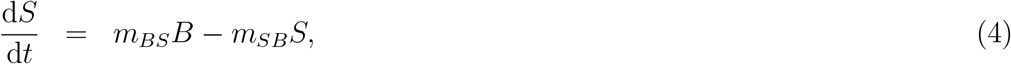

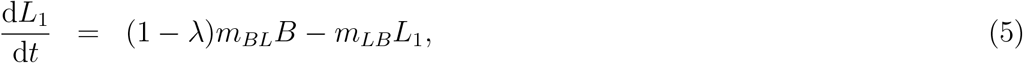

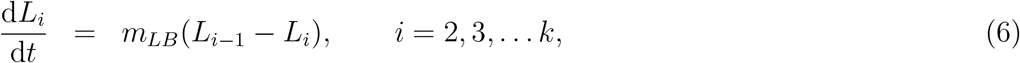

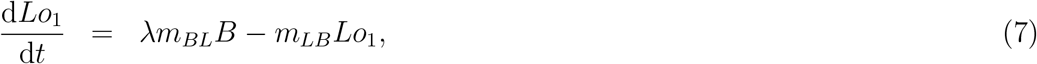

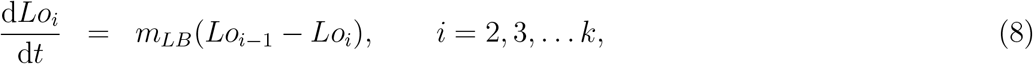

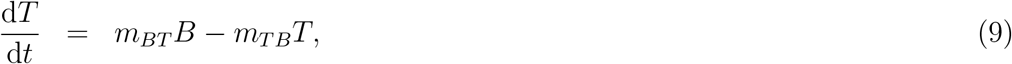

where cells exiting the cannulated lymph node, *m*_*LB*_*Lo*_*k*_ do not return to the blood because they are sampled, and *m*_*LB*_*Lo*_*k*_ is the rate of labeled lymphocyte exit from the sampled lymph node (which is compared to experimental data, e.g., column 2 in the dataset #1, and see Figure 2C&D).

Because in our experimental data, measurements of labeled lymphocytes were done either in the blood and efferent lymph (or only efferent lymph), the assumption of *n* = 3 tissue compartments (spleen, LNs, and other non-lymphoid tissues) may not be always justified. Furthermore, the number of sub-compartments *k* in LNs is also unknown. Therefore, in our analyses we fitted a series of models assuming different values for *k* and *n* to the data from Frost *et al.* [39] (dataset #1) and compared the quality of the model fit to data using AIC (see Results section).

The average residence time of lymphocytes in the spleen or non-lymphoid tissues is 1*/m*_*SB*_ and 1*/m*_*T B*_, respectively. The residence time of lymphocytes in the LNs is given as *RT* = *k/m*_*LB*_. The initial number of labeled cells in the blood varied by experiment and is indicated in individual graphs. In some fits parameter *λ* could not be identified from the data and thus was fixed (indicated by absent predicted confidence intervals). The rest of the parameters were fit by the model.

##### Blood-LN dynamics model

Many studies of lymphocyte recirculation that report kinetics of accumulation and loss of labeled lymphocytes in efferent lymph of ovine LNs do not report dynamics of these cells in the blood which is the major limitation of such studies. Therefore, to gain insight into whether LN cannulation data alone can be used to infer lymphocyte residence times in the LNs we propose an alternative model which only considers lymphocyte dynamics in the blood, one LN, and the efferent lymph of the cannulated LN. In this “blood-LN dynamics” model (Figure 1C) the dynamics of lymphocytes in the blood is given by a phenomenological function *B* given as a sum of *j* declining exponentials. The rationale to use such a model stems from the kinetics of labeled lymphocyte dynamics in the blood observed in Frost *et al.* [39] and studies (Figure 2A&B and [41]) even though actual dynamics in other experimental systems may not follow the same pattern. The dynamics of labeled cells in the sampled lymph node is thus driven by the continuous entry of labeled cells from the blood into the LN. The model is then given by following equations

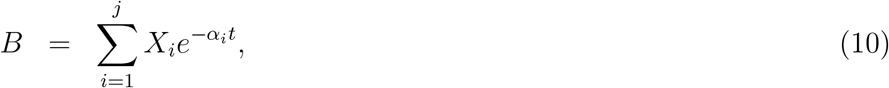

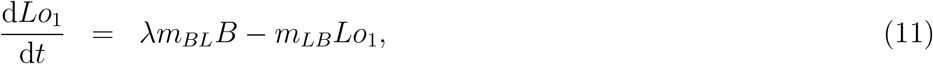

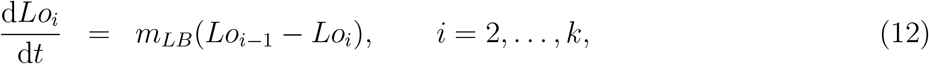

*X*_*i*_ and *α*_*i*_ are the initial values and the rate of decline in the *i*^*th*^ exponential function, *λm*_*BL*_ is the overall rate at which lymphocytes from the blood enter the LN, and *m*_*LB*_ is the rate at which lymphocytes move between *k* sub-compartments in the LN and exit the LN. The rate at which lymphocytes exit the LN and thus are sampled in the LN efferent lymph is *m*_*LB*_*Lo*_*k*_. It should be noted that parameters *λ*, *m*_*BL*_ and *X*_*i*_ in many cases are not identified from the data on lymphocyte dynamics in the efferent lymph, thus, the main parameter that we are interested in is the average residence time of lymphocytes in the LNs given by *RT* = *k/m*_*LB*_. Because the dynamics of cells in the blood is generally unknown when fitting the model predictions to data, we varied the number of exponential functions *j* = 1, 2, 3 and compared the quality of fits of different models using AIC.

#### 2.2.3 Migration when cells are injected into afferent lymph

In one set of experiments migration of labeled lymphocytes via LNs was measured by directly injecting lymphocytes into the afferent lymph of a LN and observing accumulation and loss of these cells in the efferent lymph of the LN. To use these data to estimate the lymphocyte residence time in the LN we assume that cells injected into afferent lymph *A* migrate into the lymph node at a rate *m*_*A*_, and then the cells migrate via each of *k* sub-compartments in the LN at a rate *m*_*LB*_. With these assumptions the dynamics of cells in the afferent lymph and the LN are given by equations:

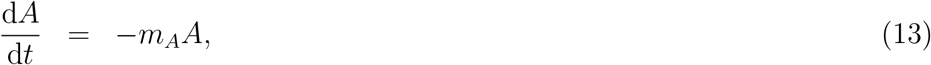

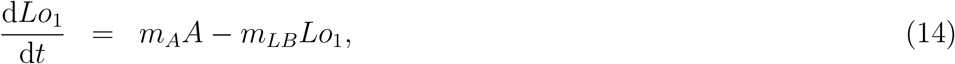

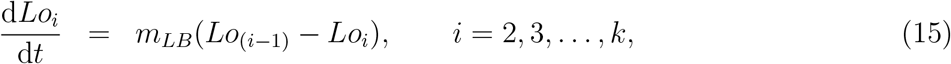

where initially all labeled cells were in the afferent lymph. As previously stated, the rate of lymphocyte exit from the LN via efferent lymph is given by *m*_*LB*_*Lo*_*k*_. The average residence time of lymphocytes in the LN is then *RT* = *k*/*m*_*LB*_.

#### 2.2.4 Homing to different lymph nodes

In the final set of experiments Reynolds *et al.* [20] collected lymphocytes from efferent lymph of pLN or mLN, labeled and then re-infused the collected cells i.v. into the same animal. The labeled cells were then collected in the efferent lymph of the pLN and mLN. Because the authors did not report the dynamics of labeled cells in the blood, we extended the ‘blood-LN dynamics’ model (see eqns. (10)–(12)) to describe cell migration from the blood to the efferent lymph of two LNs. The number of labeled lymphocytes found in the *i*^*th*^ sub-compartment of the pLN and mLN are given by *Lo*_1*,i*_ and *Lo*_2*,i*_, respectively.

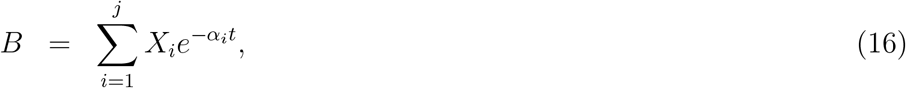

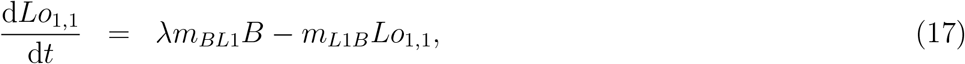

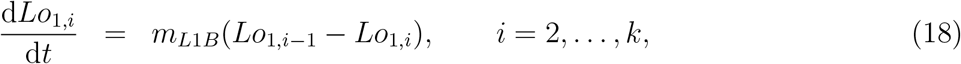

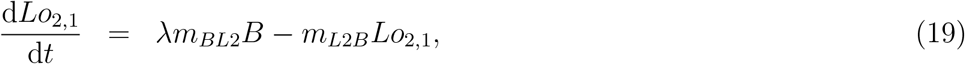

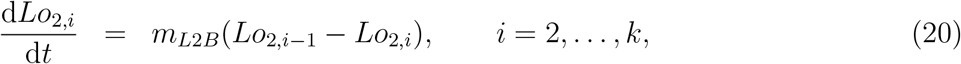

where *m*_*L*1*B*_ and *m*_*L*2*B*_ are the rate of lymphocyte exit from the pLN and mLN, respectively, *m*_*BL*1_ and *m*_*BL*2_ are the rates of lymphocyte entry from the blood to pLN and mLN, respectively, and *j* = 1, 2 in fitting models to data. Because the data clearly showed the difference in accumulation of lymphocytes in different LNs, we considered two alternative explanation for this difference. In one model we assume that the difference in kinetics is due to differences in the rate of lymphocyte entry into specific LNs (*m*_*BL*1_ ≠ *m*_*BL*2_) while residence times are identical in the two LNs (*m*_L1*B*_ = *m*_*L*2*B*_). In the alternative model, the rate of entry into the LNs are the same but residence times may differ (*m*_*BL*1_ = *m*_*BL*2_ and *m*_*L*1*B*_ ≠ *m*_*L*2*B*_).

#### 2.2.5 Statistics

The models were fitted to data in R (version 3.1.0) using **`modFit`** routine in **`FME`** package by logtransforming the data (single or two different measurements) and model predictions and by minimizing the sum of squared residuals. Numerical solutions to the system of equations were obtained using ODE solver lsoda (from the deSolve package) with default absolute and relative error tolerance. Different algorithms such as `BFGS`, `L-BFGS-B`, or **`Marquart`** in the **`modFit`** routine were used to find parameter estimates. Discrimination between alternative models was done using corrected Akaike Information Criterion, AIC [50]

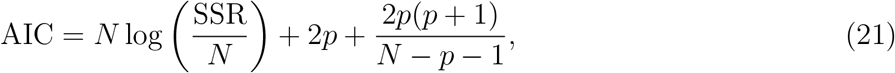

where *SSR* is the sum of squared residuals, *N* is the number of data points, and *p* is the number of model parameters fitted to data. The model with the minimal AIC score among all tested models was viewed as the best fit model, but a difference of AIC score of 1 − 3 between best fit and second best fit models was generally viewed as not significant [50]. Predicted 95% confidence intervals for estimated parameters were calculated as ±2*σ* with standard deviation *σ* provided for each parameter by the **`modFit`** routine.

## 3 Results

### 3.1 Estimating lymphocyte residency time in the LNs using lymphocytes dynamics in efferent lymph

To study recirculation of different subsets of lymphocytes in sheep, typical experiments involve isolation of lymphocytes from a tissue (e.g., blood, LN, efferent lymph of a LN), labeling of the isolated lymphocytes, re-infusion of the labeled cells into the same animal, and monitoring of the concentration of labeled cells in different tissues, typically the efferent lymph of LNs [35, Figure 1A]. In one such experiment, Frost *et al.* [39] collected lymphocytes from the efferent lymph of a pLN, labeled the cells with ^51^Cr and re-injected the cells into the sheep i.v. The authors measured concentration of labeled cells in the blood or the percent of labeled cells that were exiting a pLN [39]. We scaled these data and calculated the total number of labeled cells in the blood (Figure 2A) or the number of labeled cells exiting the pLN per unit of time (Figure 2C, see Materials and Methods for more detail). To describe the dynamics of the labeled cells in the blood and lymph simultaneously, we developed two alternative mathematical models: recirculation model and blood-LN dynamics model (Figure 1B&C and see Materials and Methods for more detail). The recirculation model assumes that migration of lymphocytes occurs between major secondary lymphoid tissues, while the blood-LN dynamics model uses phenomenologically described dynamics of labeled cells in the blood to predict the dynamics of labeled cells in the efferent lymph of the LN. Both models were first fitted to the experimental data on labeled lymphocyte dynamics in the blood and efferent lymph in the first 90 hours after cell transfer (“short-term migration” data).

While the overall structure of the recirculation model was defined by the number *n* of different compartments through which lymphocytes could recirculate (Figure 1B), we investigated how many such compartments are in fact needed to describe the experimental data. Therefore, we fitted a series of recirculation models with a variable number of compartments and determined how well such models described the data. In addition, we also tested how many sub-compartments *k* in the LNs are needed for best description of the data (see eqns. (3)–(9)). The analysis revealed that *n* = 3 tissue compartments and *k* = 3 sub-compartments in the LNs are needed to adequately describe the dynamics of labeled cells in the blood and efferent lymph (Table 1). Such a model could accurately describe simultaneously the loss of labeled cells in the blood and accumulation and loss of labeled cells in the efferent lymph (Figure 2A&C). The model predicted existence of two recirculation compartments with the average residence times of 2.4 and 19.5 hours, with the latter compartment corresponding to LNs in the sheep. The nature of the first compartment is unclear but given the estimated residence time it is likely that the first compartment represent spleen (e.g., see [43]). The final third compartment was needed to explain the long-term loss of labeled cells from the blood at a rate of about *d* = 0.02/h. The predicted rate of lymphocyte migration to tissues (*m*_*BS*_ + *m*_*BL*_ + *m*_*BT*_ ≈ 1.4 − 1.6/h ≫ *d*, see Table 2) is higher than the observed rate *d* because of the return of lymphocytes that had migrated to lymphoid tissues back to circulation. The model also naturally explains the long-term decline in the number of labeled lymphocytes found in the efferent lymph of the cannulated pLN which is simply driven by the decline of labeled cells in the blood. The analysis also suggests that none of the obvious characteristics of the distribution of the lymphocyte exit rate from the LN such as the time of the peak or the average of the overall distribution (e.g., see Figure 2C) accurately represent the average residence time. This result strongly suggests that to accurately estimate lymphocyte residence times from LN cannulation experiments it is critical to use appropriate mathematical models.

**Table 1:**
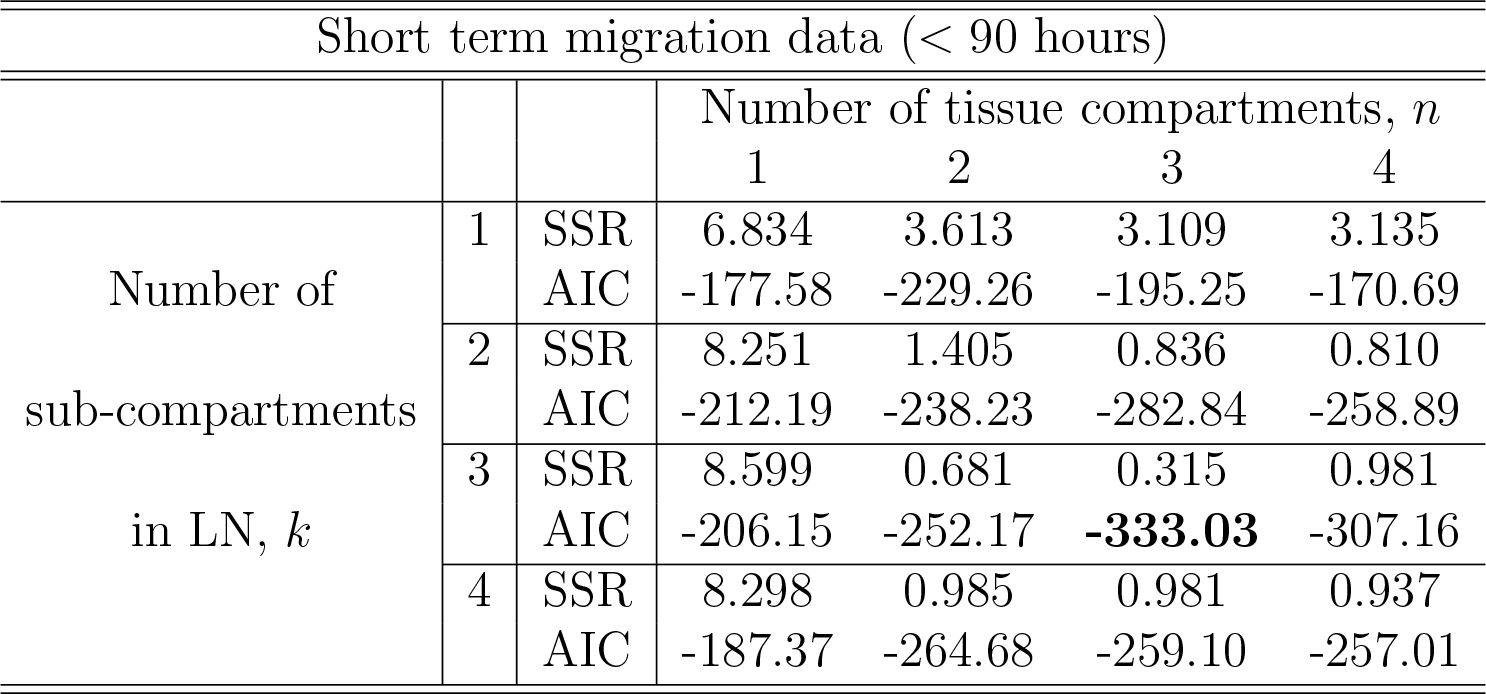
Comparison of different recirculation mathematical models fitted to the data on RL dynamics in blood and prescapular LN. Experiments were performed as described in Figures 1 and 2 and a series of mathematical models assuming recirculation of lymphocytes via *n* different tissue compartments with LNs having *k* sub-compartments (eqns. (3)–(9)) were fitted to experimental data (shown in Figure 2A&C). We tested *n* = 1 … 4 different tissue compartments with *k* = 1 … 4 sub-compartments in LNs. The bold AIC value shows the model of best fit with *n* = 3 and *k* = 3. Parameters for the best fit model are shown in Table 2 and the best fit is shown in Figure 2A&C.

**Table 2:**
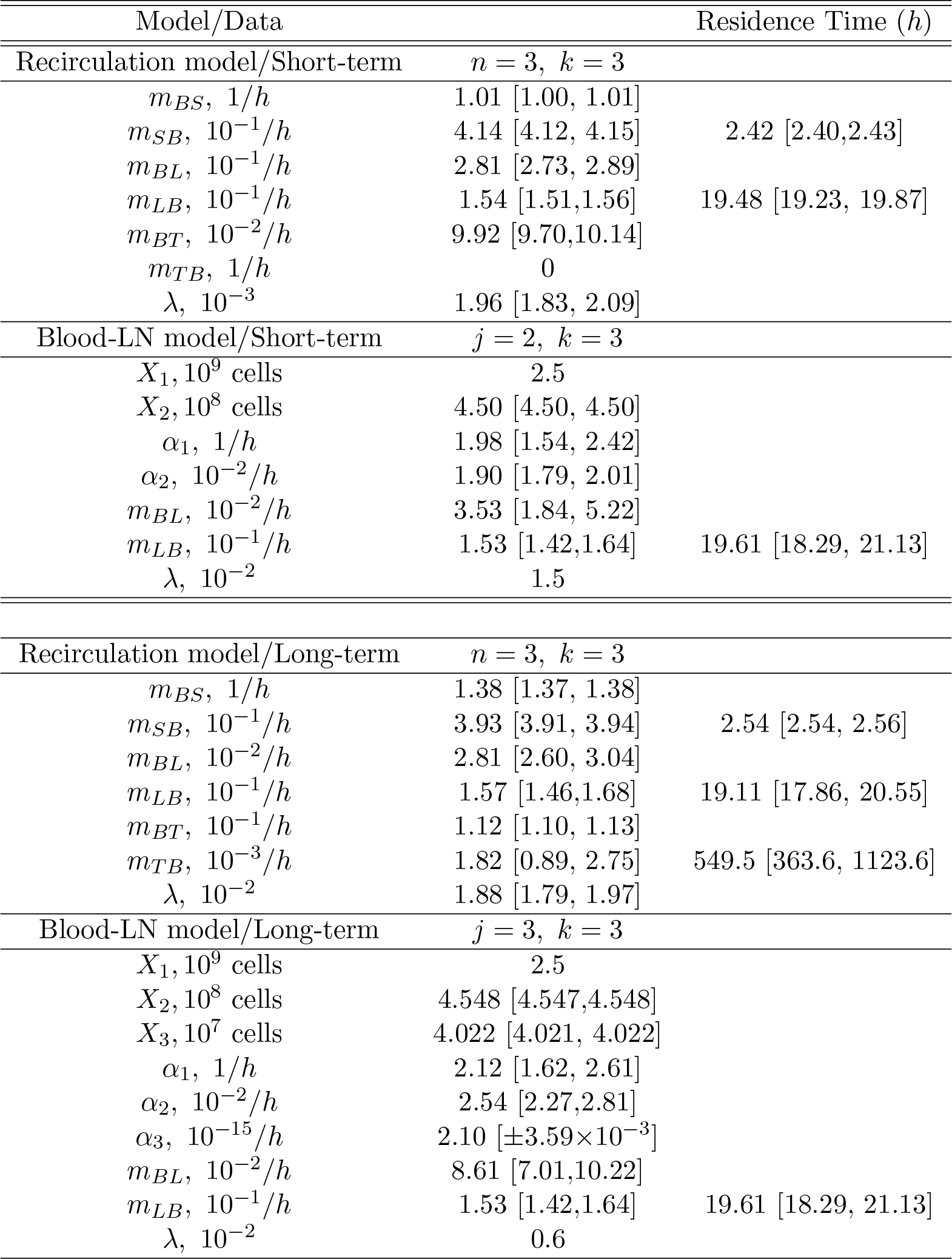
Parameters of the best fit recirculation model (eqns. (3)–(9)) or the blood-LN dynamics model (eqns. (10)–(12)) fitted to either short-term migration data (*t <* 90 *h*) or the long-term migration data of Frost *et al.* [39] (see Materials and Methods for more detail). In the recirculation model the three tissue compartments are suggested to be spleen, LNs, and other non-lymphoid tissues and the migration rates from the blood to these compartments are denoted as *mij* with *i, j* = *B, S, L, T*. In the blood-LN dynamics model it was not possible to estimate accurately the initial number of labeled lymphocytes in the blood (*X*_1_), so that parameter was fixed to *X*_1_ = 2.5 × 109 cells. Residence times in LNs were calculated as *RT* = *k/mLB* and as *miB* for other compartments (*i* = *S, T*).

Assuming a smaller (*k* = 1) or a larger (*k* = 4) number of sub-compartments in the LNs resulted in poorer fits of the data (Table 1). The intuitive reason of why the model in which lymphocyte residence times in LNs are exponentially distributed (*k* = 1) does not fit the data well follows from the rapid loss of labeled lymphocytes in the blood within the first hours after lymphocyte transfer (Figure 2A). Rapid decline in the number of labeled lymphocytes in the blood reduces the rate at which new labeled cells enter the pLN which would have resulted in a relatively rapid exit of cells from the pLN for exponentially distributed residence time. Similarly, the model in which there are too many sub-compartments would force the distribution of cells in the efferent lymph to be even broader, thus, also resulting in poorer fit. Thus, this analysis suggests that migration of lymphocytes via LNs is not described by a simple exponential function and there is a requirement for lymphocytes to spend some minimal time in LNs before exiting into circulation.

It is interesting to note how the dynamics of labeled lymphocytes in the blood may be used to infer recirculation kinetics of cells. Indeed, the initial rapid decline of the number of labeled lymphocytes is explained in the model by migration to secondary lymphoid tissues and change in the decline rate at 2-3 hours after lymphocyte transfer is naturally explained by the exit of initially migrated cells from one of the compartments (most likely spleen) back to the blood. Thus, lymphocyte kinetics in the blood suggests residence time in first compartment of about 2-3 hours (Table 2).

The recirculation model makes a strong assumption that the dynamics of labeled lymphocytes in the blood and efferent lymph are due to migration of lymphocytes into and out of different tissues (Figure 1B). Without measurement of lymphocyte accumulation and loss in other secondary lymphoid tissues such as spleen, predictions of the recirculation model remain speculative. Therefore, to estimate residence times of lymphocytes in LNs we developed an alternative mathematical model which links the lymphocyte dynamics in efferent lymph to cell dynamics in the blood. This model involves a small number of assumptions, the major of which is the physiological constraint that most lymphocytes enter lymph nodes from from blood [2]. Intuitively, however, this model allows us to estimate the time taken by cells to migrate from the blood to the efferent lymph of a LN irrespective of the specific route of this migration.

In the model the dynamics of labeled lymphocytes in the blood is described phenomenologically as a sum of several exponential functions, and by fitting a series of such models we found that the dynamics of labeled cells in the first 90 hours after cell transfer is best described by a sum of two exponentials (Table S1 and Figure 2B). The model predicted a rapid initial loss of lymphocytes in the blood at a rate of 2/h (half-life time of about 21 min) and a slower loss rate of 0.02/h after the first 4 hours (half-life time of 35 hours, Figure 2B and Table 2).

By fitting a series of mathematical models in which the number of sub-compartments in the pLN was varied, to the data on dynamics of labeled lymphocytes in the efferent lymph we found that *k* = 3 sub-compartments provided fits of the best quality (Table S1 and Figure 2D). Importantly, the model predicted the average residence time of lymphocytes in the pLN of 19.6 h which is nearly identical to the value found by fitting recirculation model to the same data (Table 2).

Both recirculation and blood-LN dynamics models were then fitted to the long-term migration data in which the dynamics of labeled cells in efferent lymph was measured continuously for 120 hours while in the blood there was an extra measurement at 120 hours (Figure 3A&B). The recirculation model with *n* = 3 tissue compartments and *k* = 3 sub-compartments was able to accurately describe the data although the model underestimated the number of labeled lymphocytes in the blood at 120 hours post-transfer (Figure 3A and Table S2). Interestingly, the model required a non-zero rate of lymphocyte return from the third tissue back to circulation (Figure 3A and Table 2). The latter requirement stemmed from the fact that in the absence of lymphocyte return from the last tissue compartment the model would match worse the number of labeled cells in the blood (results not shown). Importantly, the recirculation model predicted similar average residence times of lymphocytes in the first two compartments (representing spleen and LNs) to that of the model fitted to short-term dataset (Table 2).

Perhaps unsurprisingly, to describe the dynamics of labeled lymphocytes in the blood over 120 hours the sum of three different exponential functions was required (Table S1). Furthermore, the model with *k* = 3 sub-compartments in the pLN was able to describe the dynamics of labeled lymphocytes in efferent lymph with best quality (Table S1 and Figure 3B&D). The model predicted the average residence time of lymphocytes in LNs to be 19.6 h which is consistent with results from the recirculation model fitted to the same data or the models fitted to short-term migration data.

Taken together, analysis of data from Frost *et al.* [39] on recirculation of lymphocytes via prescapular LN in sheep suggests non-exponentially distributed residence times of lymphocytes in the LNs with the average time being approximately 20 h. Developed mathematical models naturally explain the long-term presence of labeled lymphocytes in efferent lymph node of a cannulated LN by continuous input of new labeled cells from the blood to the LN.

### 3.2 Migration of lymphocytes from afferent to efferent lymph suggests non-exponentially distributed residence time in LN

Analysis on the dynamics of labeled lymphocytes transferred i.v. into sheep suggested that migration of lymphocytes via LN follows a multi-stage process which can be described as cell migration via identical sub-compartments (Figure 1B&C). Since the time it takes for lymphocytes to cross the endothelial barrier and enter LNs is very short (few minutes, [51]), the finding that distribution of lymphocyte residence times are not exponential could still be due to some unknown processes.

Therefore, to further investigate the issue of the distribution of residence times of lymphocytes in ovine LNs we analyzed another experimental dataset. In these experiments, Young *et al.* [47] isolated lymphocytes from the efferent lymph of the pLN, labeled and injected the cells into the afferent lymph of the popliteal LN (poLN), and then measured exit of the labeled cells the efferent lymph of the poLN (Figure 4 and see Materials and Methods for more detail). Cells, injected into the afferent lymph, cannot move to any other tissue but the draining LN, and thus, such data allow to directly evaluate kinetics of lymphocyte migration via individual LN.

To describe these experimental data, we adapted the blood-LN dynamics model to include migration of labeled lymphocytes from the afferent lymph to the LN and then to the efferent lymph (eqns. (13)–(15)). The model has 3 unknown parameters that must be estimated from the data (*A*(0), *m*_*A*_, *m*_*LB*_). Unfortunately, the original data for cell dynamics for individual animals were not available, and the digitize data only included 3 time points which did not allow accurate estimation of all model parameters (results not shown). Therefore, to investigate the dynamics of labeled cells in the efferent lymph we fitted a series of mathematical models with a varying number of sub-compartments *k* in the LN and average residence times *RT* = *k/m*_*LB*_ fixed to several different values to the experimental data (Table S3). Analysis revealed that several sub-compartments are needed for accurate description of the data and the actual number of sub-compartments varied for different cell subtypes, but was never less than *k* = 3 (Table S3). The expected residence times also varied with the cell type but overall were within 18-20 hour range which is consistent with the previous analysis of Frost *et al.* [39] data (Figure 4).

By fitting the data with the model in which the number of sub-compartments *k* was varied we found that the estimated residence time of lymphocytes in the poLN was dependent on the assumed *k* (Table S4). This is consistent with our recent result on estimating residence time of T and B lymphocytes in LNs of mice using the data from photoconvertable Kaede mice [9]. Interestingly, the model fit predicted a relatively slow movement of lymphocytes from the afferent lymph to the LN, which is determined by the parameter *m*_*A*_ (1*/m*_*A*_ ≈ 5 h) and was dependent on the number of subcompartments. These results also support our conclusion that residence times of lymphocytes in ovine poLN are not exponentially distributed and the average residence time for different lymphocyte subsets is around 20 hours.

### 3.3 Impact of lymphocyte kinetics in the blood on estimates of the lymphocyte residency time in the LNs

Our analysis of the Frost *et al.* [39] data demonstrated the usefulness of having measurements of the dynamics of labeled lymphocytes both in the blood and efferent lymph of a specific cannulated LN. Unfortunately, many of the published studies that we have surveyed lacked measurements of lymphocyte counts in the blood and only recorded cell numbers in the efferent lymph, often as the percent of labeled cells in the overall population. An important question is whether the estimates of the residency time of lymphocytes in LNs, found by fitting mathematical models to the data on lymphocyte counts in efferent lymph, depend on the assumed lymphocyte dynamics in the blood.

In several different experiments the decline of i.v. injected labeled lymphocytes in the blood is bi-exponential with the rapid decline in cell numbers within a few hours and slower decline in the next days [41, see Figure 2B]. By fitting the blood-LN dynamics model (with *k* = 3) to the data on the dynamics of labeled lymphocytes in efferent lymph (shown in Figure 2D) we found that fits of similar quality could be obtained independently whether the dynamics of labeled cells in the blood follow either a single or bi-exponential decline (results not shown). However, the estimates of the average residence time of lymphocytes in LNs was dependent on the assumed model of lymphocyte dynamics in the blood; namely, assuming a bi-exponential decline resulted in longer average residence times (results not shown).

To investigate this issue further we fitted the blood-LN dynamics model (with *k* = 3) assuming that the number of labeled cells in the blood follows an exponential decline, to the data on labeled cell dynamics in efferent lymph. In this analyses we either fitted the rate of cell decline in the blood (*α*_1_) or fixed it to different values (Figure 5). We found that the decline rate of labeled cells in the blood has a dramatic impact on the quality of the model fit of the data as well as on estimates of the average residence times (Figure 5B). In particular, assuming that the number of labeled cells remains constant in the blood (*α*_1_ = 0) predicts a constant output of labeled cells in efferent lymph, and as the result, failed to accurately describe the data (Figure 5B). Similarly, assuming that the loss of labeled cells occurs relatively rapidly (*α*_1_ = 0.05/*h*) also results in poor fits of the data and longer average residence time of lymphocytes in the LN (Figure 5B). However, allowing the rate of lymphocyte loss *α*_1_ to be fitted allowed the model describe the data well further suggesting that the 4 long-term dynamics of labeled cells in the efferent lymph is the consequence of cell dynamics in the blood.

**Figure 5:**
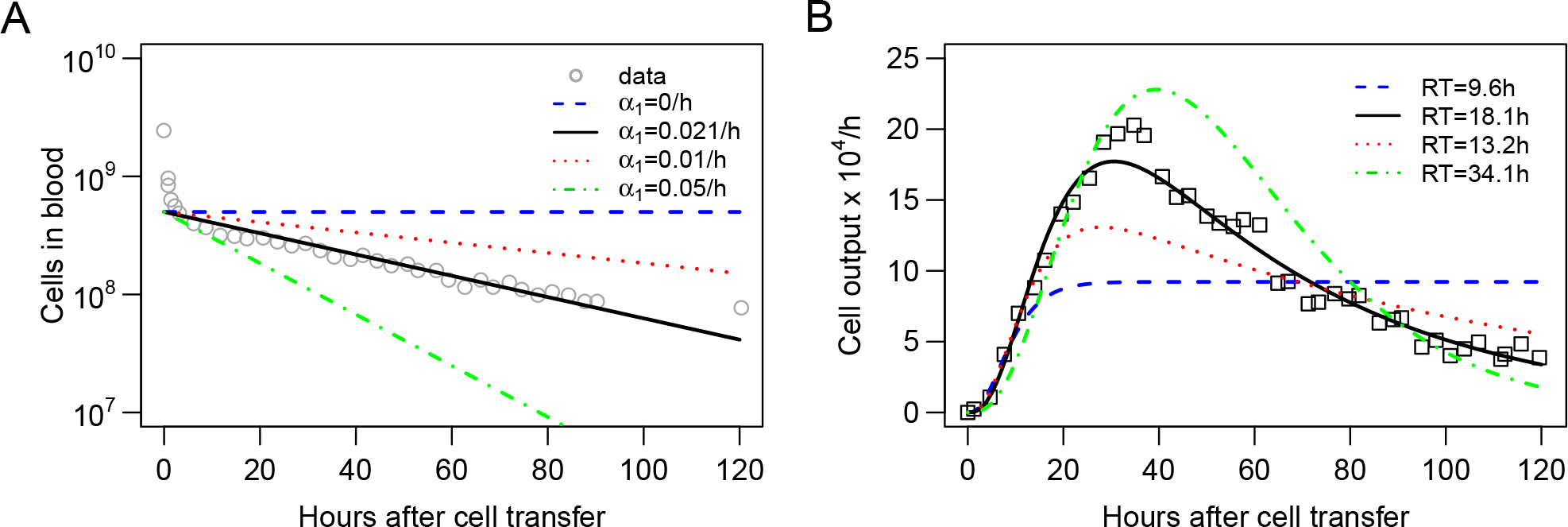
Kinetics of lymphocyte loss in the blood influences the estimate of the lymphocyte residence time in LNs. We fit the blood-LN dynamics model (eqns. (10)–(12)) to the data on accumulation and loss of labeled lymphocytes in the efferent lymph of the pLN of sheep (shown by markers in panel B; see Materials and Methods for details on the data) assuming that loss of the lymphocytes in the blood occurs exponentially at a rate *α*_1_ (panel A). In contrast with other analyses, fits of the model to data in this case were done without log10 transformation. In the fits the decline rate *α*_1_ was either fixed to several different values (*α*_1_ = 0, *α*_1_ = 0.01*/h*, or *α*_1_ = 0.05*/h*) or was estimated by fitting the model to data in panel B (*α*_1_ = 0.021*/h*). The estimated residence time of lymphocytes in the LN *RT* is indicated in panel B. In the fits we assumed that LNs have *k* = 3 sub-compartments. The initial number of labeled lymphocytes in the blood *X*_1_ = 5 × 108 and the proportion of lymphocytes migrating to the prescapular LN *λ* = 0.01 were fixed in model fits of the data. In panel A the data on labeled lymphocyte dynamics in the blood from Frost *et al.* [39] was not used in model fitting and is only displayed by markers for illustrative purposes.

### 3.4 Lymphocytes migrate more rapidly to LNs which the cells recently exited

All data analyzed so far have been for lymphocytes isolated from prescapular LNs which migrate back to prescapular or popliteal LNs. An important question is whether the average residence time of lymphocytes varies across types of LN. Indeed, previous analysis of migration of naive T cells in mice suggest that lymphocytes spend less time in gut-draining mesenteric LNs than in skin-draining LNs [42]. In contrast, another study suggested similar residency times of thoracic duct lymphocytes in skin- and gut-draining LNs of rats [43]. To address this issue we analyzed another experimental dataset.

In their studies, Reynolds *et al.* [20] isolated T lymphocytes from efferent lymph of prescapular or mesenteric LNs (pLN and mLN, respectively), labeled them with different fluorescent dyes, reinjected the cells into the same animal, and measured accumulation and loss of the labeled cells in efferent lymph of pLN and mLN (Figure 6 and see Materials and Methods for more detail). The data showed that T cells isolated from pLN accumulate to higher numbers in the efferent lymph of pLN as compared to cells from mLN and vice versa (Figure 7). There could be at least two alternative explanations for such differential accumulation of cells in LNs of their origin: preferential migration or preferential retention. According to the preferential migration hypothesis, cells from pLN have a higher rate of entry into pLN than the rate at which cells from mLN enter the pLN (and vice versa). In contrast, in the preferential retention hypothesis, cells from pLN have a longer residence time in pLN as compared to cells from mLN (and vice versa).

**Figure 6:**
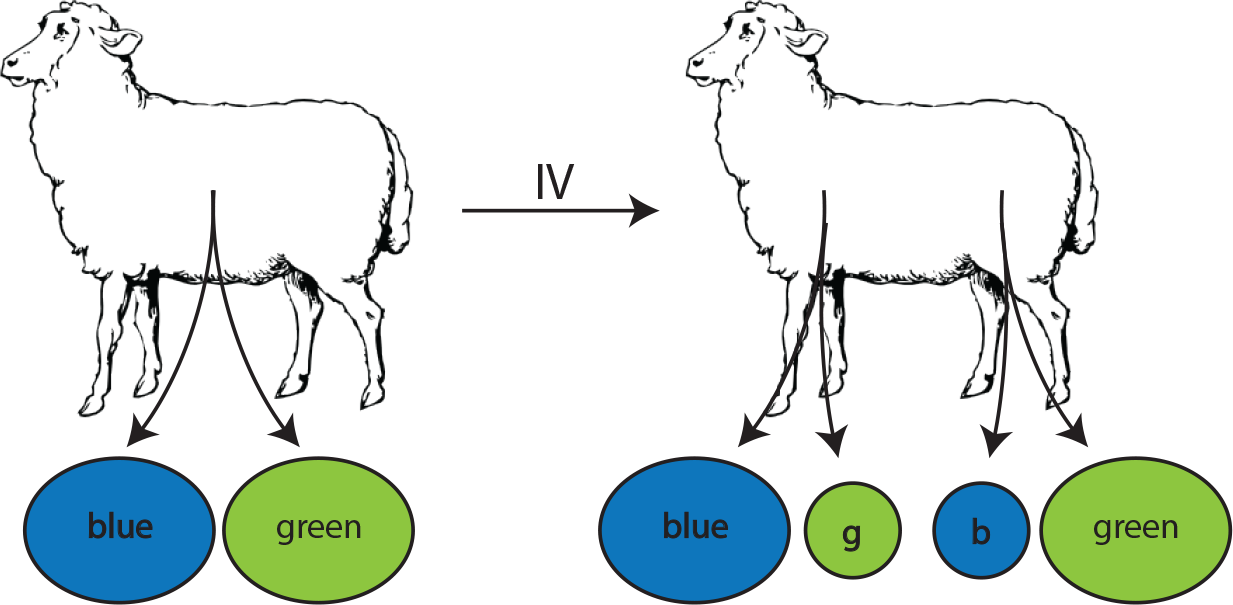
Experimental design of the study to evaluate migration of lymphocytes isolated from different LNs. Populations of lymphocytes were isolated from efferent lymph by cannulation of either the pLN (skin-draining) or the mesenteric (intestine-draining) lymph node (mLN), labeled with RITC (green) or FITC (blue), respectively. These cells were then injected i.v. into the same animal. Accumulation and loss of the injected lymphocytes was followed in efferent lymph of pLN and mLN over time.

**Figure 7:**
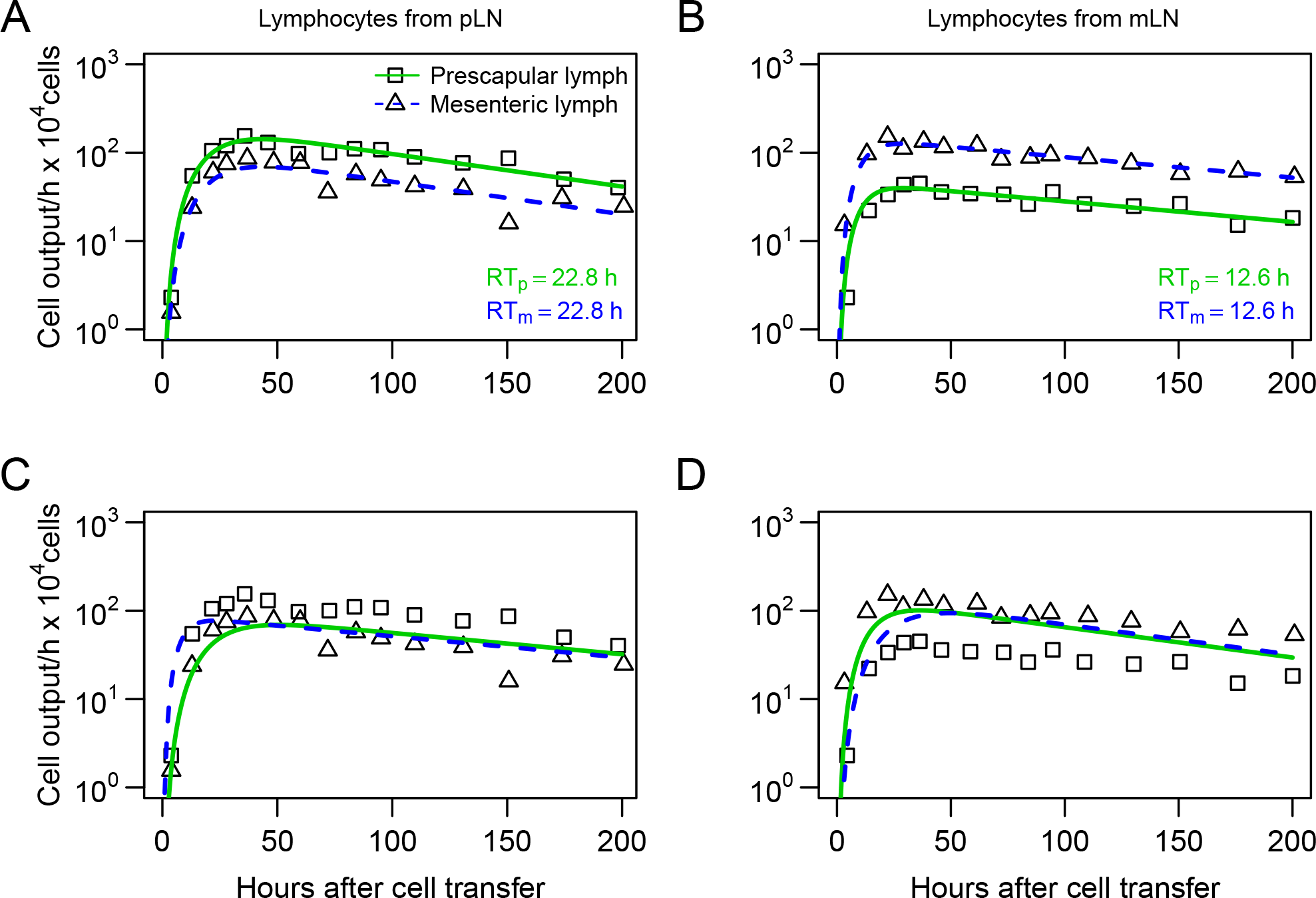
Lymphocytes have a higher entry rate into the LN which they recently exited. Experiments were performed as described in Figure 6 and the number of cells exiting pLN (boxes) or mLN (triangles) was recorded for lymphocytes collected previous from pLN (panels A&C) or from mLN (panels B&D). We fitted mathematical model (eqns. (16)–(20)) to the cannulation data either assuming that difference in lymphocyte dynamics is due to differential entry of lymphocytes into pLN and mLN (panels A&B) or is due to differential residence time of lymphocytes in the LNs (panels C&D). Markers show the data and lines are predictions of the mathematical models. Model arameters are shown in Table S5. The average residence times of lymphocytes in pLN (*RT*_*p*_) and mLN (*RT*_*m*_) predicted by the best fit model are shown on panels A&B.

To discriminate between these alternative hypotheses, we fitted the blood-LN dynamics model to these data. Specifically, we assumed that the dynamics of labeled T cells in the blood follow bi-exponential decline and that lymphocytes must traverse via *k* = 3 sub-compartments in the LNs.

In the preferential migration model we fixed the average residence times of lymphocytes in LNs for cells from pLN and mLN (determined by the parameter *m*_*LB*_) but allowed different rates of entry into the LN from the blood (determined by the parameter *m*_*BL*_, see eqns. (16)–(20)). This model could accurately describe experimental data (Figure 7A&B). Interestingly, the model predicted 2 fold higher entry rate into pLN by cells of pLN origin and 3 fold higher entry rate into mLN by cells of mLN origin as compared to cells of pLN origin (Table S5). Importantly, assuming identical average residence time of T cells from pLN in skin-draining or gut-draining LNs resulted in fits of excellent quality suggesting the average residence time of T cells from pLN does not depend on the LN type. However, T cells from mLN migrated via LNs nearly 2 fold faster than T cells from pLN suggesting that the average residence time does depend on the origin of T cells.

In the alternative preferential retention model we fixed the rate of lymphocyte entry from the blood to the LNs and allowed the residence times (or more precisely, the rate of exit of T cells from the LNs) to vary depending on the LN type. This model failed to accurately describe the data (Figure 7C&D) suggesting that the data cannot be explained solely by increased retention of cells in the LN of their origin. Importantly, allowing both entry and exit rates to depend on the LN type did not improve the model fit of the data for lymphocytes from pLN (*F*_1,24_ = 0.26, *p* = 0.62) but marginally improved the fit of the data for lymphocytes from mLN (*F*_1,24_ = 6.3, *p* = 0.02).

## 4 Discussion

It is well understood that some lymphocytes are able to recirculate between blood and secondary lymphoid tissues such as lymph nodes. In part, this understanding came from multiple experiments on lymphocyte migration from the blood to efferent lymph of various LNs in sheep. Yet, while the data on the kinetics of lymphocyte migration via individual LNs have been published, quantitative interpretation of these data has been lacking until recently. In particular, the average residence time of lymphocytes in the ovine LNs remained largely unknown and there was incomplete understanding of why labeled lymphocytes persisted in the efferent lymph of cannulated LNs.

The first attempt known to us to explain lymphocyte dynamics in efferent lymph during cannulation experiments in sheep was by Thomas *et al.* [41] who modelled lymphocyte migration via the LN as a random walk. The model suggested that long-term detection of labeled lymphocytes in efferent lymph was due to inability of some lymphocytes to exit the LN. Here we formulated several alternative mathematical models, based on the basic understanding of lymphocyte recirculation in mammals which accurately explain the cannulation data, and proposed an alternative explanation for long term detection of labeled cells in efferent lymph. Namely, because labeled cells persist in the blood, continuous entry of such cells into the LN can naturally explain long-term persistence of labeled cells in the efferent lymph.

Our mathematical modeling approach allowed to provide novel estimates of lymphocyte residence time in ovine lymph nodes which vary between 12 and 20 hours depending on lymphocyte type and being approximately independent of the type of LN (e.g., skin- or gut-draining, Figures 2 and 7). Furthermore, the combination of data and mathematical model predicted an existence of a compartment with a shorter residence time, about 2-3 hours, which we propose is likely to be the spleen. Indeed, we recently published a similar estimate of lymphocyte residence time in the spleen of rats [43]. Parameter estimates also suggest a relatively short residence time of lymphocytes in the blood (e.g., for long-term migration data 1/(*m*_*BS*_ + *m*_*BL*_ + *m*_*BT*_) ≈ 1.6/h or *RT*_*B*_ = 28 min). This is relatively similar to previous observations [9, 43].

Another important conclusion from our analyses is that the data on lymphocyte dynamics in efferent lymph is not described well by a model in which residence times of lymphocytes in LNs are exponentially distributed (Table 1). In part, this is because of the wide distribution in the exit rates of labeled lymphocytes in efferent lymph over time. However, describing cell migration via LNs as a simple one directional process and ignoring the ability of lymphocytes to remain in the LN for longer (e.g., by including a “backflow” in cell movement as was done by Thomas *et al.* [41]) may be an over-simplification. Yet, because the model in which lymphocyte residence times are gamma distributed describes the experimental data with acceptable quality (e.g., see Figure 2), introducing additional details/parameters contradicts the fundamental “Occam’s razor” principle.

Our analysis suggests potential limitations of interpreting data from ovine LN cannulation experiments in which the dynamics of transferred lymphocytes is not tracked in the blood. In particular, we found that estimates of lymphocyte residence times in LNs do depend on the assumed model for lymphocyte dynamics in the blood (e.g., single vs. double exponential decline and slow or rapid decline). Therefore, future studies on lymphocyte recirculation kinetics in sheep should always attempt to measure and report concentration of transferred cells in the blood.

One of the fundamental questions of lymphocyte recirculation is whether lymphocytes in the blood have some “memory” of the specific LN they recently came from, and if such memory exists, whether it comes from preferential entry into a specific LN or from preferential retention in the LN. Several experimental studies have addressed the question qualitatively. For example, activated lymphocytes, or lymphoblasts, collected from the intestinal lymph of sheep were shown to accumulate preferentially in tissues associated with the gut [52], and a similar finding was reported for lymphoblasts isolated from intestinal lymph of rats [53, 54]. In contrast, lymphoblasts isolated from peripheral lymph preferentially accumulated in peripheral lymph nodes [53]. There has also been a distinction in migratory preference based on cellular subset as it has been observed that small lymphocytes accumulate in mucosal sites such as Peyer’s patches [55, 56].

We used mathematical modeling to investigate whether preferential accumulation of lymphocytes in the LN of their origin is due to preferential entry or preferential retention for one specific dataset [20]. Our analysis suggested that a model with preferential retention was not able to accurately describe the experimental data, while the model in which cells could preferentially enter a LN was able to describe the data well (Figure 7). Intuitively, this may be because the earliest increase in the number of cells found in the efferent lymph seems to be driven by rate of cell entry into the node and the data clearly indicate difference in cell accumulation in the efferent lymph depending on the cell’s origin.

There are a number of limitations with experimental data and our modeling analyses that need to be highlighted. In particular, in all of our experimental data, the dynamics of labeled lymphocytes in the efferent lymph was reported as a frequency of total cells, which required recalculation to determine the total number of cells exiting a specific LN per unit of time (e.g., [39]). Similarly, calculation of the total number of lymphocytes in the blood requires the knowledge of the total blood volume of animals which was not reported. The required recalculations may introduce errors (e.g., due to incorrectly assumed blood volume in the animals) and thus may influence the values for some estimated model parameters. For example, a smaller assumed blood volume in animals would naturally lead to a lower number of transferred lymphocytes in Frost *et al.* [39] experiments detected in the blood which should directly impact the estimate of the rate of lymphocyte migration from the blood to the LN. In fact, the absolute values of estimated rates at which lymphocytes are predicted to migrate to LNs from the blood should be treated as approximate.

One of the major assumptions we made in the models was that all labeled cells have identical migratory characteristics, e.g., all cells are capable of entering and exiting the LNs and do so at the same rates. In many previous studies the types of lymphocytes used in recirculation experiments (e.g., naive or memory lymphocytes, B or T cells) were not specified, and it is very possible that migratory properties vary by cell type (e.g., see [9]). It is clear however that including multiple cell subpopulations will increase complexity of the models, making them perhaps unidentifiable from the data we have available. Also, the models based on kinetically homogeneous cell populations could describe the data reasonably well, which suggests that there is no need to introduce a more complex models. Yet, comparison of predictions found by the best fit models for the data did indicate some discrepancy, for example, the best fit model was not fully capable of capturing the peak of the exit rate of labeled lymphocytes in efferent lymph (e.g., Figure 5B). It is possible that including slow and fast recirculating cell sub-populations may be able to fully capture the peak in labeled cells even though we were not able to improve fit of these data by extending the model to two sub-populations with different migration kinetics (results not shown).

Another major assumption of our modeling approach is that lymphocytes in circulation enter LNs via HEVs and not via afferent lymph of the tissues. While there is some experimental support for this assumption for lymphocytes migrating in non-inflammatory conditions [2], it is clear that during inflammation in the skin, many cells may enter the skin-draining LNs via afferent lymph [19].

At its core, the combination of experimental data and our models allowed us to estimate the time it takes for lymphocytes to migrate from the blood to the efferent lymph of specific LNs — and our results suggest that this time is gamma distributed. For lymphocytes migrating to LNs via HEVs, this distribution is likely to be related to lymphocyte residence time in LNs as lymphocytes pass via HEV rather quickly [51]. However, if lymphocytes migrate from the blood to efferent lymph by first entering non-lymphoid tissues (e.g., skin), then exiting the tissue into afferent lymphatics, and then pass via the LN — then our estimates of the average residence time of lymphocytes in LNs are upper bound values.

In most of our models we ignored the possibility of cell death. When describing labeled lymphocyte dynamics during short-term (< 90 h) migration experiments with the recirculation model (Figure 2) we found the need to have a tissue compartment which acts as a sink and thus may represent a death process (Table 2). However, there appears to be an equilibrium reached by recirculating lymphocytes in the blood by 120 h of the experiment (Figure 3A) suggesting a limited role of death process in determining overall dynamics of labeled lymphocytes. Still, we performed some additional analyses by adding death rate to all tissue compartments and found that the best fit is found when such death rate is small or non-existent (Table S6).

Even with all limitations in the data and assumptions of the models, we provided a quantitative framework to analyze data from LN cannulation experiments in sheep. The models, developed in the paper, may need to be tailored to explain kinetics of lymphocyte recirculation in specific experiments. As illustrated in this work, greater insights into mechanisms regulating lymphocyte migration in large animals such as sheep and humans may thus be obtained by combining the use of quantitative experiments and mathematical modeling.

## Author’s contributions

The study was originally designed by VVG. Data were digitized from cited publications by MMD. All major analyses were performed by MMD. The paper was written by MMD and VVG.

## Acknowledgments

We would like to thank the immunology community for discussion over this research, especially Michio Tomura, Gudrun Debes, and David Masopust. This work was in part supported by the NIH grant (R01 GM118553) to VVG.

