## Supplemental Information for "Estimating residence times of lymphocytes in ovine lymph nodes"

### Supplemental Material

| Short-term migration data ($t < 90$ $h$) | | | | | |
|---|---|---|---|---|---|
| | | | Number of exponentials $j$ | | |
| | | | 1 | 2 | 3 |
| # of SC in LN, $k$ | 1 | SSR | 6.421 | 1.256 | 1.256 |
| | | AIC | -141.80 | -243.09 | -237.97 |
| | 2 | SSR | 5.409 | 0.379 | 0.278 |
| | | AIC | -152.94 | -320.95 | -336.02 |
| | 3 | SSR | 6.389 | 0.295 | 0.249 |
| | | AIC | -142.12 | **-337. 26** | -335.29 |
| | 4 | SSR | 8.224 | 0.897 | 0.784 |
| | | AIC | -125.71 | -264.94 | -268.57 |

| Long-term migration data | | | | |
|---|---|---|---|---|
| | | Number of exponentials $j$ | | |
| | | 1 | 2 | 3 |
| 1 | SSR | 7.310 | 1.483 | 1.483 |
| | AIC | -169.39 | -285.97 | -281.04 |
| 2 | SSR | 6.393 | 0.517 | 0.371 |
| | AIC | -179.58 | -366.10 | -386.10 |
| 3 | SSR | 8.020 | 0.384 | 0.349 |
| | AIC | -162.35 | -388.63 | **-391.09** |
| 4 | SSR | 10.354 | 0.959 | 0.855 |
| | AIC | -142.94 | -319.06 | -322.93 |

**Supplemental Table S1:** Comparing quality of fits of different models to the data dynamics of recirculating lymphocytes in the blood and efferent lymph of prescapular LN (see Figure 2 and 3 in the main text). Quality of the fits of the mathematical model (eqns. (10)–(12)) as judged by SSR and AIC are shown for different numbers of exponential functions for the cell dynamics in the blood ($j = 1 \ldots 3$) or the different number of sub-compartments (SC) in the LN ($k = 1 \ldots 4$). The bold values indicate lowest AIC for the best fit models.

| Long-term migration T → LN → B | | | | |
|---|---|---|---|---|
| | | | Number of Comp. $n$ | |
| | | | 3 | 4 |
| # of SC in LN ($k$) | 2 | SSR | 1.072 | 0.948 |
| | | AIC | -305.73 | -309.80 |
| | 3 | SSR | 0.657 | 0.647 |
| | | AIC | **-359.81** | -338.84 |
| | 4 | SSR | 0.982 | 0.981 |
| | | AIC | -312.35 | -307.21 |

| Long-term migration T → B | | | | |
|---|---|---|---|---|
| | | | Number of Comp. $n$ | |
| | | | 3 | 4 |
| # of SC in LN ($k$) | 2 | SSR | 0.535 | 0.575 |
| | | AIC | -361.04 | -318.56 |
| | 3 | SSR | 0.574 | 0.545 |
| | | AIC | **-384.13** | -354.50 |
| | 4 | SSR | 0.824 | 0.721 |
| | | AIC | -314.75 | -334.45 |

**Supplemental Table S2:** Long term lymphocyte migration is best modeled using a recirculation model when with $n = 3$ tissue compartments and $k = 3$ sub-compartments in the LNs, and lymphocytes exiting non-lymphoid tissues return directly to the blood instead of draining into lymph nodes first. The SSR and AIC values for models fitted to the long term data are shown as $n$ and $k$ are altered. The left table shows results of fitting the recirculation model in which there is a migration of lymphocytes from non-lymphoid tissues to LNs at a rate $\rho$. The right table shows results of the basic recirculation model in which lymphocytes return to the blood directly when exiting the non-lymphoid tissue. Both data from LN output and decay in blood were used for fitting the models. The bold AIC values shows the best fit model.

| *RT* | | $k=2$ | $k=3$ | $k=4$ | $k=5$ | $k=6$ | $k=7$ | $k=8$ |
|---|---|---|---|---|---|---|---|---|
| 15 h | CD4 T cells | 11.086 | 9.804 | 8.505 | 7.212 | 5.891 | 4.520 | **3.074** |
| | CD8 T cells | 10.075 | 10.285 | 7.516 | 6.355 | 5.294 | 4.006 | **2.791** |
| | $\gamma\delta$ T cells | 11.798 | 12.398 | 6.913 | 4.676 | 2.062 | -1.230 | **-6.002** |
| | B cells | 11.019 | 10.503 | 4.231 | 3.406 | 6.811 | **-0.571** | -0.433 |
| 17.5 h | CD4 T cells | 10.741 | 8.979 | 6.858 | 4.212 | 0.664 | -4.949 | **-24.291** |
| | CD8 T cells | 9.504 | 7.832 | 5.894 | 3.608 | 0.807 | -2.887 | **-8.661** |
| | $\gamma\delta$ T cells | 10.293 | 7.604 | 3.885 | -2.332 | **-25.438** | -2.403 | 2.425 |
| | B cells | 6.686 | 4.578 | 2.031 | -1.204 | -5.786 | -14.658 | **-16.926** |
| 20 h | CD4 T cells | 10.989 | 8.497 | 5.793 | 1.568 | -6.406 | **-11.096** | -1.139 |
| | CD8 T cells | 10.226 | 8.664 | 4.535 | 0.863 | -4.939 | **-24.999** | -7.916 |
| | $\gamma\delta$ T cells | 12.697 | 6.924 | 1.116 | **-22.714** | -0.555 | 4.703 | 7.426 |
| | B cells | 9.527 | 6.987 | -0.078 | -6.105 | **-33.855** | -8.081 | -3.129 |
| 22.5 h | CD4 T cells | 10.516 | 8.326 | 5.299 | 0.552 | **-11.152** | -5.300 | 1.523 |
| | CD8 T cells | 9.092 | 6.762 | 3.549 | -1.414 | **-13.029** | -8.474 | -1.936 |
| | $\gamma\delta$ T cells | 9.871 | 6.249 | -0.453 | **-11.231** | 2.418 | 6.795 | 9.188 |
| | B cells | 6.073 | 2.930 | -1.915 | **-12.788** | -9.823 | -2.918 | 0.087 |
| 25 h | CD4 T cells | 10.562 | 9.917 | 6.103 | 1.926 | -4.628 | **-20.871** | -1.413 |
| | CD8 T cells | 10.303 | 6.581 | 3.839 | -2.474 | **-22.139** | -5.338 | -0.188 |
| | $\gamma\delta$ T cells | 10.889 | 9.329 | 0.112 | **-18.225** | 5.854 | 5.666 | 8.304 |
| | B cells | 9.972 | 4.722 | -3.254 | **-24.551** | -5.983 | -0.963 | 1.476 |

**Supplemental Table S3:** Determining best model describing migration of lymphocytes from the afferent to efferent lymph (Young *et al.* [47] data). We fitted mathematical model (eqns. (13)–(15) in main text) to the data of lymphocyte accumulation and loss in efferent lymph (see Figure 4 in the main text) by fixing the number of sub-compartments in the LN and fixing the rate of lymphocyte exit from each sub-compartment. Obtained AIC values are listed in the table. Bolded black values show the value for the best fit per cell type and residence time, while bolded red values show overall best fit per cell type.

| # of SC | Cell type | $A$, % | $m_A$, $1/hr$ | $m_{LB}$, $1/h$ | $RT$, $h$ | SSR |
|---|---|---|---|---|---|---|
| $k = 5$ | CD4 T cells | 16.15 | 0.224 | 0.224 | 22.32 | 8.391e-02 |
| | CD8 T cells | 14.49 | 0.199 | 0.198 | 25.25 | 3.941e-02 |
| | $\gamma\delta$ T cells | 20.09 | 0.427 | 0.195 | 25.64 | 4.437e-29 |
| | B cells | 14.29 | 0.151 | 0.188 | 26.59 | 6.903e-31 |
| $k = 6$ | CD4 T cells | 18.55 | 0.269 | 0.269 | 22.30 | 4.418e-03 |
| | CD8 T cells | 16.41 | 0.238 | 0.238 | 25.21 | 2.831e-04 |
| | $\gamma\delta$ T cells | 37.63 | 0.080 | 0.345 | 17.39 | 3.195e-29 |
| | B cells | 26.72 | 0.053 | 0.299 | 20.06 | 6.952e-30 |
| $k = 7$ | CD4 T cells | 25.43 | 0.140 | 0.368 | 19.02 | 2.268e-30 |
| | CD8 T cells | 25.10 | 0.094 | 0.346 | 20.23 | 2.639e-29 |
| | $\gamma\delta$ T cells | 49.06 | 0.067 | 0.431 | 16.25 | 9.348e-29 |
| | B cells | 40.42 | 0.036 | 0.386 | 18.14 | 1.233e-30 |
| $k = 8$ | CD4 T cells | 32.26 | 0.109 | 0.454 | 17.62 | 1.775e-29 |
| | CD8 T cells | 33.34 | 0.070 | 0.431 | 18.56 | 2.707e-29 |
| | $\gamma\delta$ T cells | 60.30 | 0.060 | 0.514 | 15.56 | 1.056e-28 |
| | B cells | 55.98 | 0.028 | 0.468 | 17.09 | 1.145e-30 |

**Supplemental Table S4:** Evaluating impact of the number of assumed sub-compartments in the LNs on the estimate of the average residence time of lymphocytes in LNs. We fitted mathematical model (eqns. (13)–(15) in the main text) to the data of lymphocyte accumulation and loss in efferent lymph (see Figure 4 in the main text) by fixing the number of sub-compartments per lymph nodes ($k$) and by fitting the rate of lymphocyte entry into the LN ($m_{A1}$), lymphocyte exit from the LN ($m_{1A}$), and the initial percent of labeled cells in the afferent lymph ($A$). Low SSR values indicate that some versions of the model fit the data perfectly suggesting over-fitting.

| Prescapular Origin | Prescapular LN ($i = 1$) | | Mesenteric LN ($i = 2$) | |
|---|---|---|---|---|
| Parameter | Fixed Res. Time | Fixed Entry | Fixed Res. Time | Fixed Entry |
| $m_{BLi},\ 10^{-2}/h$ | 3.70 [2.99, 4.41] | 1.70 [1.01, 2.39] | 1.80 [1.45, 2.15] | 1.70 [1.01, 2.39] |
| $m_{LiB},\ 10^{-1}/h$ | 1.32 [1.13,1.51] | 1.21 [0.49, 1.92] | 1.32 [1.13,1.51] | 3.11 [1.40, 4.81] |
| $\alpha_2,\ 10^{-3}/h$ | 8.49 [6.48,10.40] | 5.56 [1.23, 9.88] | 8.49 [6.48,10.40] | 5.56 [1.23, 9.88] |
| $RT,\ h$ | 22.8 [19.9, 26.6] | 24.8 [15.6, 61.5] | 22.8 [19.9, 26.6] | 9.7 [6.2, 21.4] |
| AIC | **-133.63** | -77.76 | **-133.63** | -77.76 |
| SSR | 0.253 | 1.631 | 0.253 | 1.631 |

| Mesenteric Origin | Prescapular LN ($i = 1$) | | Mesenteric LN ($i = 2$) | |
|---|---|---|---|---|
| Parameter | Fixed Res. Time | Fixed Entry | Fixed Res. Time | Fixed Entry |
| $m_{BLi},\ 10^{-2}/h$ | 0.90 [0.81, 0.99] | 2.48 [1.68, 3.29] | 2.84 [2.54, 3.15] | 2.48 [1.68, 3.29] |
| $m_{LiB},\ 10^{-1}/h$ | 2.38 [2.11, 2.65] | 1.67 [0.83, 2.52] | 2.38 [2.11, 2.65] | 1.14 [0.84, 1.44] |
| $\alpha_2,\ 10^{-3}/h$ | 5.35 [4.38, 6.32] | 7.90 [4.20, 11.61] | 5.35 [4.38, 6.32] | 7.90 [4.20, 11.61] |
| $RT,\ h$ | 12.6 [11.3, 14.2] | 18.0 [11.9, 36.2] | 12.6 [11.3, 14.2] | 26.4 [20.8, 35.7] |
| AIC | **-165.91** | -97.02 | **-165.91** | -97.02 |
| SSR | 0.086 | 0.858 | 0.086 | 0.858 |

**Supplemental Table S5:** Parameter estimates of the blood-LN dynamics model (eqns. (10)–(12)) fitted to the data on lymphocyte dynamics in efferent lymph in prescapular and mesenteric LNs (see Figures 6 and 7 in the main text for more detail). The following parameters were fixed for each prediction: $X_1 = 1.5 \times 10^9$ cells , $X_2 = 5.3 \times 10^8$ cells, $\alpha_1 = 3/h$, and $\lambda = 10^{-5}$; other parameters $m_{b1}$ (exit rate from blood to LNs), $m_{1b}$ (the exit rate from each sub-compartment in LNs), and $\alpha_2$ were fitted. When fitting the model by fixing residence the rate of cells exiting each LN were set to be equal. When fitting the model by fixing entry the rate of cells entering each LN was set to be equal. The LN compartment of the model has $k = 3$ sub-compartments.

| Parameter | Default | fit $\delta$ | fix $\delta$ | fix $\delta$ |
|---|---|---|---|---|
| $m_{BS},\ 1/h$ | 1.01 | 0.86 | 1.19 | 0.34 |
| $m_{SB},\ 10^{-1}/h$ | 4.14 | 5.20 | 4.42 | 3.09 |
| $m_{BL},\ 10^{-1}/h$ | 2.81 | 5.19 | 2.27 | 5.14 |
| $m_{LB},\ 10^{-1}/h$ | 1.54 | 1.39 | 1.53 | 1.33 |
| $m_{BT},\ 10^{-2}/h$ | 9.92 | 0 | 9.52 | 4.91 |
| $\lambda,\ 10^{-3}/h$ | 1.96 | 1.43 | 2.46 | 1.31 |
| $\delta,\ 10^{-2}/h$ | 0 | 1.59 | 0.001 | 0.01 |
| AIC | -333.04 | -327.25 | **-333.84** | -315.21 |
| $RT,\ h$ | 19.47 | 21.60 | 19.57 | 22.54 |

**Supplemental Table S6:** Introducing death rate of recirculating lymphocytes does not improve the quality of the model fit to data. We fitted the recirculation model (eqns. (3)–(9) in the main text) to the data on lymphocyte dynamics in the blood and the efferent lymph from Frost *et al.* [39] (Default), and allowed for extra mortality of lymphocytes $\delta$. One alternative model assumed no migration of lymphocytes to non-lymphoid tissues ($m_{BT} = 0$) and the cells die at a rate of $\delta$ from every compartment including blood. Two alternative models consider migration of lymphocytes to non-lymphoid tissues ($m_{BT} > 0$) and fixing the rate of cell death $\delta$ to two different values; these models assumed only death in the spleen and LNs.
